# Designer small molecule control system based on Minocycline induced disruption of protein-protein interaction

**DOI:** 10.1101/2023.08.15.553207

**Authors:** Ram Jha, Alexander Kinna, Alastair Hotblack, Reyisa Bughda, Anna Bulek, Isaac Gannon, Tudor Ilca, Christopher Allen, Katarina Lamb, Abigail Dolor, Farhaan Parekh, James Sillibourne, Shaun Cordoba, Shimobi Onuoha, Simon Thomas, Mathieu Ferrari, Martin Pule

## Abstract

A versatile, safe, and effective small-molecule control system is highly desirable for clinical cell therapy applications. Therefore, we developed a two-component small-molecule control system based on the disruption of protein-protein interactions using minocycline, an FDA-approved antibiotic with wide availability, excellent bio-distribution, and low toxicity. The system comprises an anti-minocycline single-domain antibody (sdAb) and a minocycline-displaceable cyclic peptide.

Here we show how this versatile system can be applied to OFF-switch split CAR systems (MinoCAR) and universal CAR adaptors (MinoUniCAR) with reversible, transient, and dose-dependent suppression; to a tunable T cell activation module based on MyD88/CD40 signaling; to a controllable cellular payload secretion system based on IL-12 KDEL retention and as a cell/cell inducible junction.

This work represents an important step forward in the development of a remote-controlled system to precisely control the timing, intensity, and safety of therapeutic interventions.

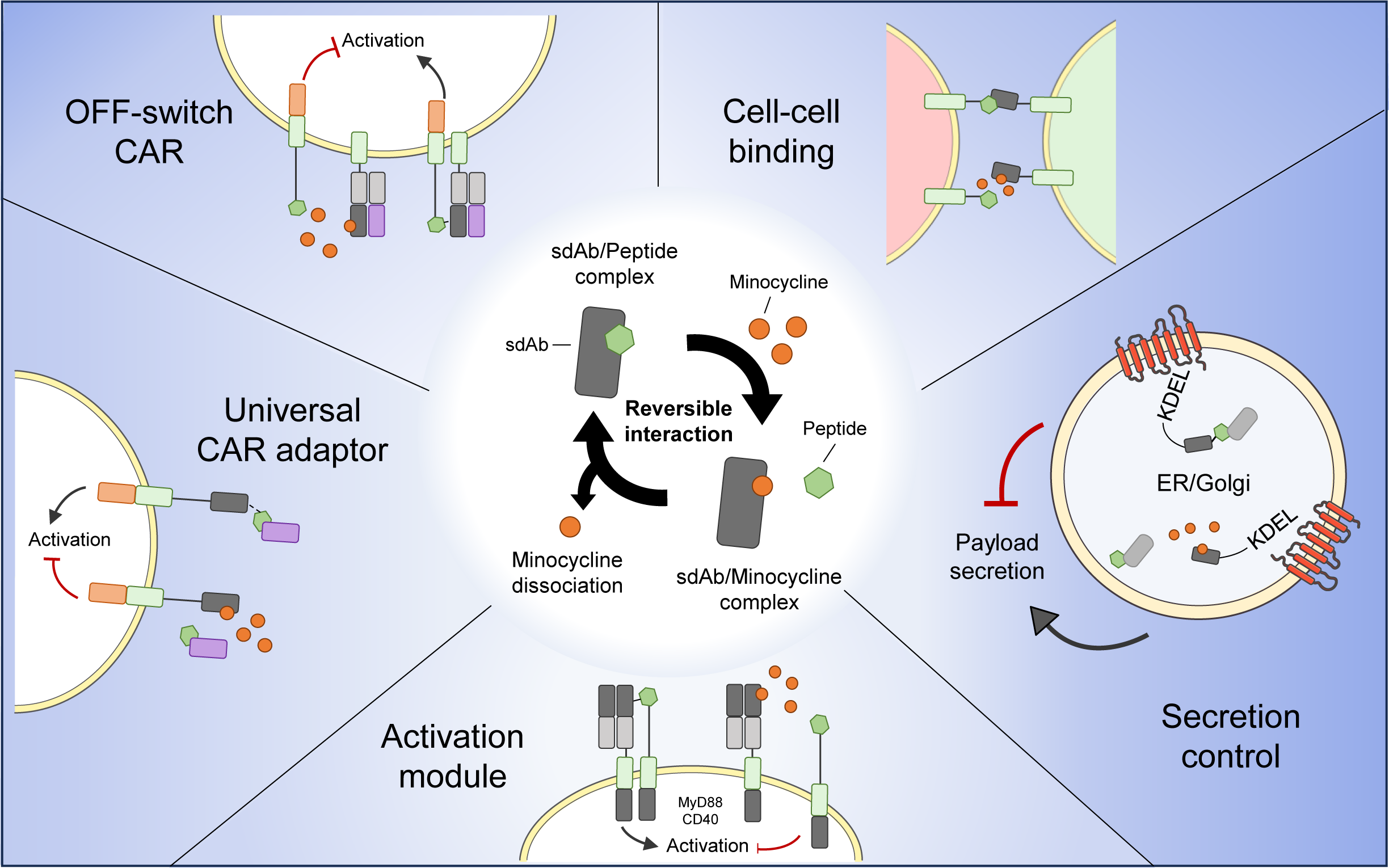

## INTRODUCTION

Engineered cellular therapies have emerged as a promising approach for treating a wide range of diseases, including cancer, autoimmune disorders, and genetic diseases. Unlike small molecules or protein therapeutics however, many cellular therapies engraft in patients and may persist and expand in an autonomous fashion. Consequently, therapeutic activity cannot be easily titrated with dose, and toxicities can be progressive and fulminant. Hence, means of controlling the activity of cellular therapies remotely, for instance through administration of small molecules is desirable.

Several small molecule cellular control systems have been developed: the best characterized exploit Rapamycin’s ability to complex simultaneously with FKBP12 and the FRB fragment of mTOR/FRAP^1^. Heterodimerization of proteins fused to FKBP12 and FRB can be induced by Rapamycin. Alternatively, homodimerization can be induced in FKBP12 fusion proteins by AP1903, a homodimer analogue of Tacrolimus. This system has been used to generate suicide genes^2,3^, inducible antigen receptors^4–6^, and inducible cytokine receptors^7^. Analogous strategies using other small molecule mediated homo/heterodimerization have been described^8^.

A different strategy to control protein-protein interaction exploits proteases which can be controlled by small molecule protease inhibitors: in one example, two protein domains are separated by a herpes C virus (HCV) protease cleavage. The two protein domains are cleaved by a co-expressed HCV protease; however, in the presence of a cognate protease inhibitor, cleavage is inhibited, and hence the two protein domains do not associate^9,10^.

Alternatively, engineered protein-protein interaction can be disrupted upon exposure to a small molecule. Such systems may be more clinically convenient since a small molecule would only need to be administered in case of toxicity. One such system was described by Giordano-Attianese et al., where the Bcl-XL and the Bcl-2 homology 3 (BH3) domain of BIM were used as the heterodimerization drivers of protein-protein interaction, prevented by two clinically tested Bcl-2 inhibitors^11^. We previously described an analogous approach: by fusing one protein to a tetracycline mimicking peptide (TiP), and a second protein fused to TetRB, exposure to tetracycline can disrupt TiP/TetRB interaction^12^.

While there are a number of small molecule control systems, many of these systems are limited by either immunogenicity (HCV, TetRB), lack of availability of the small molecule (AP1903 and non-immunosuppressive rapalogs)^11,13,14^, and unwanted pharmacologic activity of the inducing small molecule (Rapamycin). Additionally, with increasingly complex cellular engineering approaches, multiple orthogonal controls may be desired.

Here we sought to develop a new small molecule control system. We selected minocycline as the ideal inducer as it is a widely used antibiotic with few pharmacological side-effects. To avoid xenogeneic proteins, we based the system on a minocycline-recognizing single domain antibody (sdAb). We additionally identified a cyclic peptide which competed with minocycline for sdAb binding to result in a protein-protein interaction control system disrupted by minocycline. We tested several applications based on these two protein domains with control effected by minocycline-induced disruption of protein-protein interactions.

## RESULTS

### Generation of minocycline specific single-domain antibodies via phage display

Minocycline specific single domain antibodies were generated by immunization of a single alpaca with KLH-conjugated minocycline and subsequent phage display panning (Fig. 1a and Extended Data Fig.1). Seroconversion was first confirmed by ELISA (Extended Data Fig. 2a) and after 2 rounds of phage panning, enrichment was observed (Extended Data Fig. 2b).

**Figure 1.**
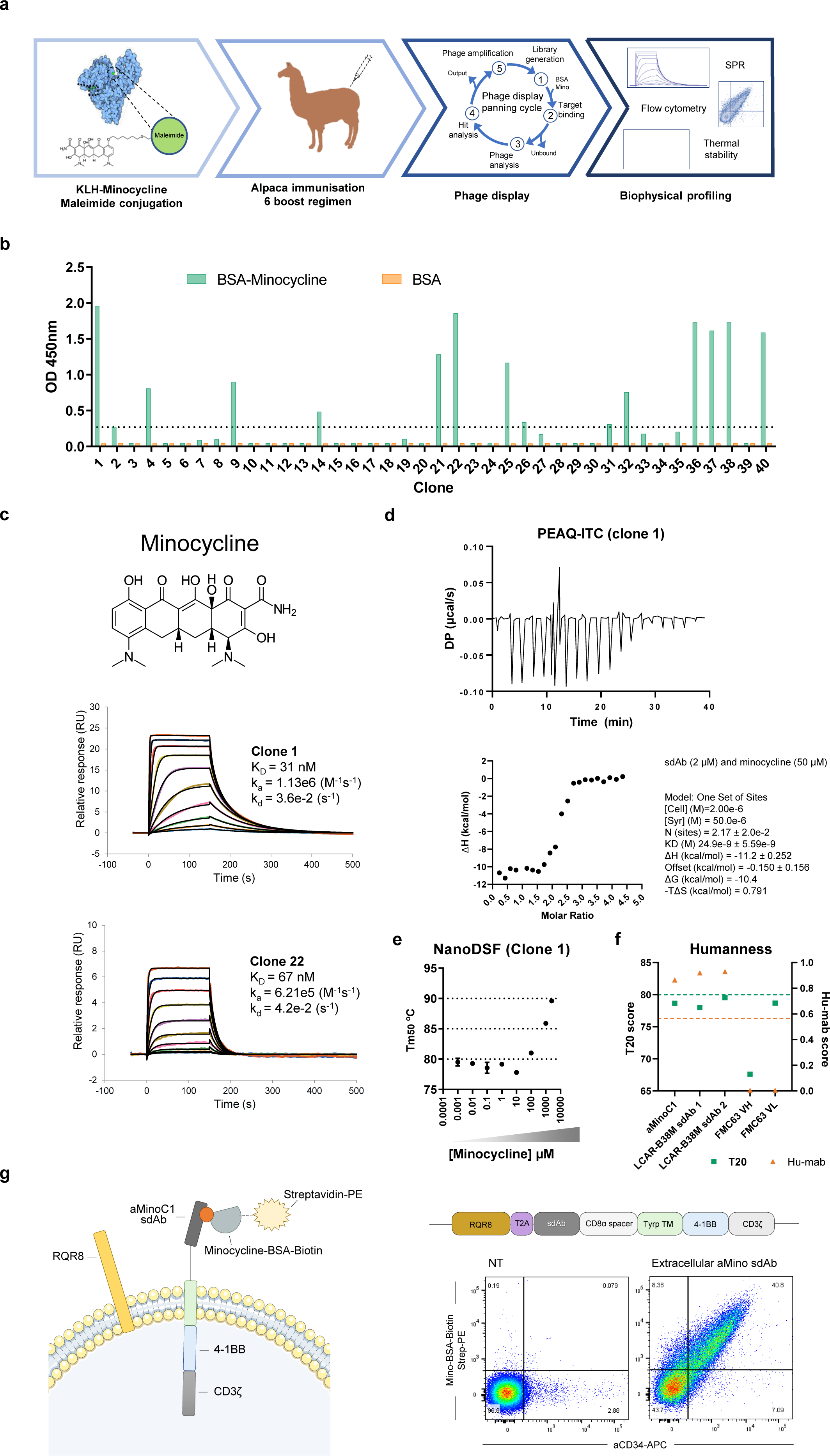
Generation and biophysical characterization of an anti-minocycline sdAb. **a**, Schematic of immunization strategy, phage display selection and biophysical characterization of anti-minocycline sdAb. **b**, ELISA screen of monoclonal anti-minocycline sdAb clones. Screening of purified monoclonal sdAb was carried out against BSA-conjugated minocycline. 14 clones showed positive binding to minocycline and a lack of binding to BSA alone control (OD > 6x baseline). **c**, Surface plasmon resonance (SPR) of anti-minocycline sdAb-Fc clone (aMinoC)1 (top) and aMinoC22 (bottom) binding to minocycline. aMinoC1 and aMinoC22 presented a KD value of 31 and 67 nM, respectively. **d**, Isothermal titration calorimetry (ITC) thermogram of aMinoC1 binding to minocycline. Raw thermogram plot over time showing ligand injection and binding saturation (top); and binding isotherm plot using one set of sites (1:1) binding model (Origin’s software) showing minocycline/aMinoC1 molar ratio and enthalpy change of the reaction (bottom) which was used to deduce a KD value of 24.9 nM (± 5.59 nM). **e**, Analysis of sdAb stability by nanoDSF. Thermal unfolding temperature (Tm) of aMinoC1 in the presence of its binding partner at concentrations from 1 nM to 1 mM. Protein concentration of 1 mg/ml suspended in PBS pH 7.4. **f**, T20 score (green) and Hu-mab score (orange) analysis of aMinoC1 sdAb in relation to LCAR-B38M sdAbs and FMC63 VH and VL domains. Dashed lines indicate threshold of human like sequences for respective scoring. **g**, Schematic representation of aMinoC1 transmembrane receptor (aMinoC1-CD8stk-TyrpTM-41BBz) and RQR8 transduction marker (left). Flow cytometry dot plot of transduced HEK293T cell surface expression for RQR8 (anti-CD34) and aMinoC1 receptor (minocycline-BSA-biotin). Linear correlation of expression between aMinoC1 receptor and RQR8 marker on cell surface.

Over 40 sdAbs from pan 2 were screened, and among the 14 clones showing specific binding to minocycline (Fig. 1b), 9 unique sequences were identified (Extended Data Fig. 2c,d). aMinoC1 showed the strongest interaction with minocycline with an SPR-determined KD of 31.6 nM (Fig. 1c). This interaction was confirmed by isothermal titration calorimetry (ITC), showing a KD of 24.9 (± 5.59) nM (Fig. 1d). aMinoC22 showed a KD of 67 nM (Fig. 1c). Neither sdAb showed an interaction with the closely related molecules doxycycline and tetracycline (Extended Data Fig. 3a).

The conformational stability of the high affinity sdAb antibody in complex with minocycline was investigated using nano differential scanning fluorimetry (nanoDSF). aMinoC1 showed high thermal stability with the first unfolding midpoint (Tm_50_) of 79.5°C. Co-incubation with minocycline ranging from 1 nM to 2.5 mM, improved the Tm_50_ values by over 10°C, rising to 89.6°C, indicating an antigen-driven stabilizing event (Fig. 1e). *In silico* analysis of aMinoC1 indicated a high humanness score, suggesting limited immunogenicity (Hu-mAb score 0.875^15^ and T20 score 78.66^16^) (Fig. 1f). Finally, the ability of aMinoC1 to be expressed on the surface of a cell and bind minocycline was shown by expressing this sdAb in a type I surface protein format and staining with labelled minocycline (Fig. 1g).

### Generation of a displaceable cyclic peptide

We next sought to identify a peptide which would compete for minocycline-dAb binding. A combinatorial phagemid library of cysteine-constrained 7-mer peptides (CX_7_C) was enriched via phage display for aMinoC1 sdAb binders. Isolated monoclonal phagemids were examined by ELISA to determine binding to sdAb and displacement by minocycline. Seventeen out of 20 selected monoclonal phage clones showed specific binding to aMinoC1 with reduction of binding when co-incubated with 1 µM minocycline (Fig. 2a). Sequencing the peptide coding region of phagemids indicated a homologous consensus motif (Pro-X-Trp-Ala-X-X-Phe) and a total of 4 unique peptide sequences with amino acid differences at positions X2, X5 and X6 (Fig. 2b).

**Figure 2.**
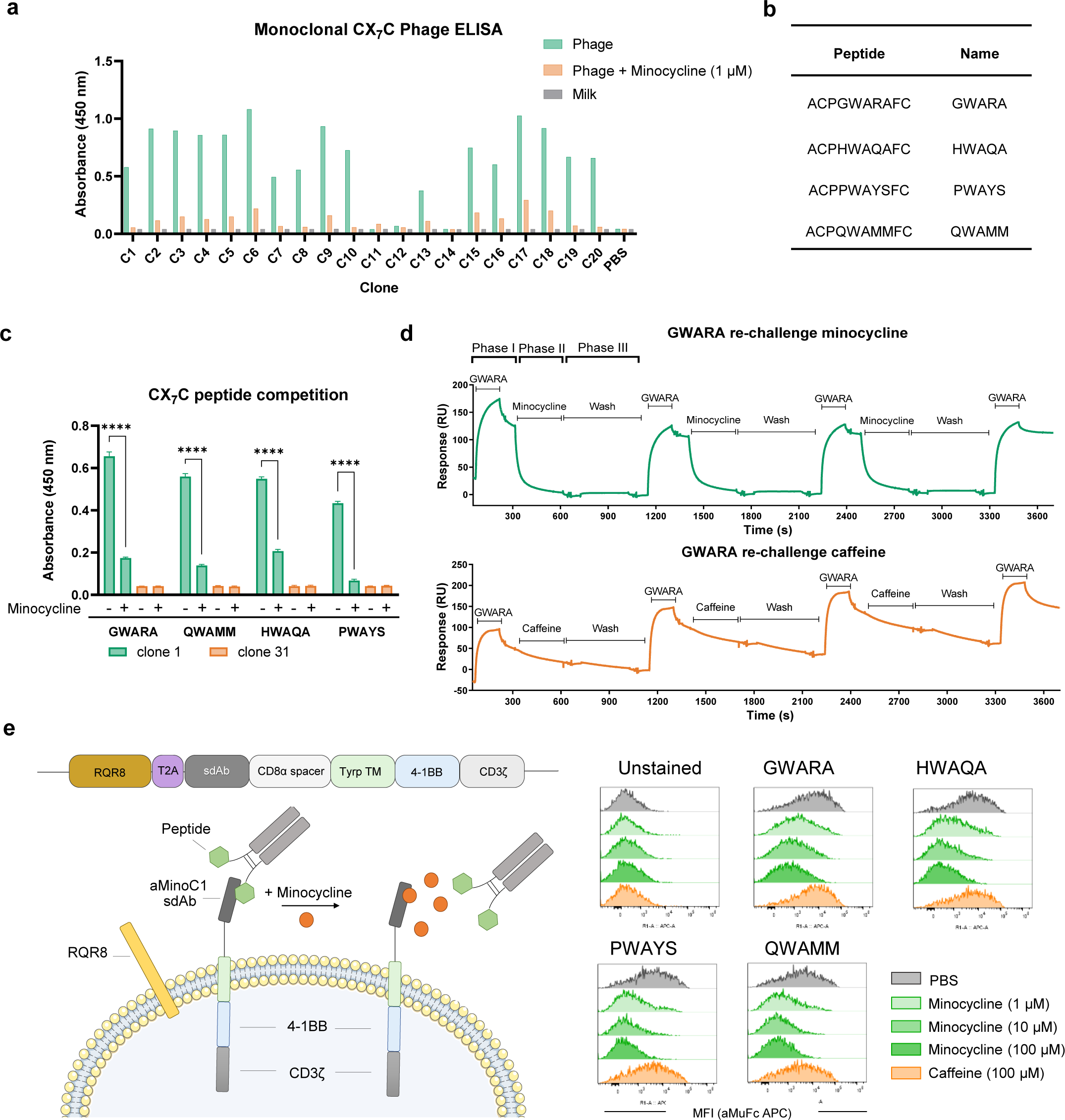
Engineering of a minocycline displaceable CX7C peptide with affinity for aMinoC1 sdAb. **a**, Anti-M13 detection ELISA of selected monoclonal whole phagemid clones displaying CX_7_C-peptides binding to plate immobilized anti-minocycline sdAb with or without minocycline (1 µM). **b**, Four unique peptide sequences (GWARA, HWAQA, PWAYS and QWAMM) were isolated and tested for sdAb binding as peptide-muIgG2a-Fc conjugates. **c**, ELISA of purified CX_7_C-muIgG2aFc peptide fusion showing specific binding to anti-minocycline sdAb. Addition of 1 µM minocycline out-competed peptide-Fc conjugates resulting in significantly reduced detection. Two-way ANOVA with Sidak’s post-test. **** P < 0.0001. **d**, Dynamic minocycline (green) or caffeine (orange) small molecules and GWARA peptide-Fc binding to immobilized aMinoC1 sdAb on Biacore T200. Sequential injections of GWARA-Fc (Phase I), small molecule (Phase II) and dissociation (buffer) step (Phase III) showing minocycline-driven acceleration of GWARA-Fc dissociation. Serial challenges with peptide and small molecule show reversibility of the system. No enhanced dissociation visible with caffeine injections. **e**, Schematic representation of cells expressing RQR8 transduction marker and aMinoC1 transmembrane receptor detected with peptide-muIgG2a Fc in the presence of minocycline (left). Histogram plot of flow cytometry staining for HEK293T transduced with aMinoC1 transmembrane receptor, with peptide-muIgG2a Fc peptide fusion (right). Dose-dependent reduction of peptide-muIgG2a Fc binding for GWARA, HWAQA, PWAYS and QWAMM in the presence of raising concentrations of minocycline (green gradient). 100 µM caffeine incubation (orange) showed no peptide binding inhibition compared to PBS control condition (grey).

The measured affinities (KD) of the GWARA, HWAQA, PWAYS and QWAMM peptides were 111 nM, 328 nM, 283 nM and 209 nM, respectively, with varying kinetic profiles, with the former showing the highest affinity and the fastest on-rate (Extended Data Fig. 3b). Specific binding of the peptides to the aMinoC1 sdAb and competition with minocycline was confirmed by ELISA as purified peptide-Fc conjugates. All peptides showed significant displacement from aMinoC1 sdAb in the presence of minocycline, while no binding was detected for the non-related sdAb clone 31 (Fig. 2c).

Reversible association and dissociation of the peptides was then demonstrated using a modified SPR protocol. Peptide binding to immobilized aMinoC1 (phase I), was specifically reversed by the addition of minocycline (phase II) and not by a non-related small molecule (caffeine). Removal of the drug (phase III) then allowed subsequent rebinding of the peptide to a comparable degree as before, confirming the sustained structural integrity of the aMinoC1 binding pocket. Serial binding/dissociation cycles confirmed robust reversibility of the system (Fig. 2d and Extended Data Fig. 4).

Minocycline was able to elicit a concentration-dependent reduction in peptide binding, for all 4 CX_7_C constructs tested, in the context of a membrane-bound aMinoC1 sdAb architecture. Additionally, incubation with caffeine at 100 µM did not result in a decrease in peptide binding (Fig. 2e and Extended Data Fig. 4).

### Understanding the minocycline and peptide binding interface on sdAb

We sought to understand the molecular interactions between the aMinoC1 sdAb and both minocycline and GWARA peptide. Crystallography failed due to low resolution of crystal formation. We subsequently performed an alanine scan, mutating all three CDR regions of the antibody and determined the effect on minocycline affinity by SPR (Fig. 3a). Mutagenesis identified a predominant role for the CDR3 region (positions 110-112 and 115-117), with additional contact points in position 38 and 55 of the CDR1 and CDR2 in binding to minocycline. Effects of alanine scanning on GWARA peptide binding were assessed by ELISA. In contrast to minocycline binding, this identified positions 28, 35 and 40 of the CDR1, positions 58, 63 and 64 of the CDR2, and position 108 of the CDR3 as the main drivers of peptide binding (Fig. 3b), suggesting a stronger role for CDRs 1 and 2.

**Figure 3.**
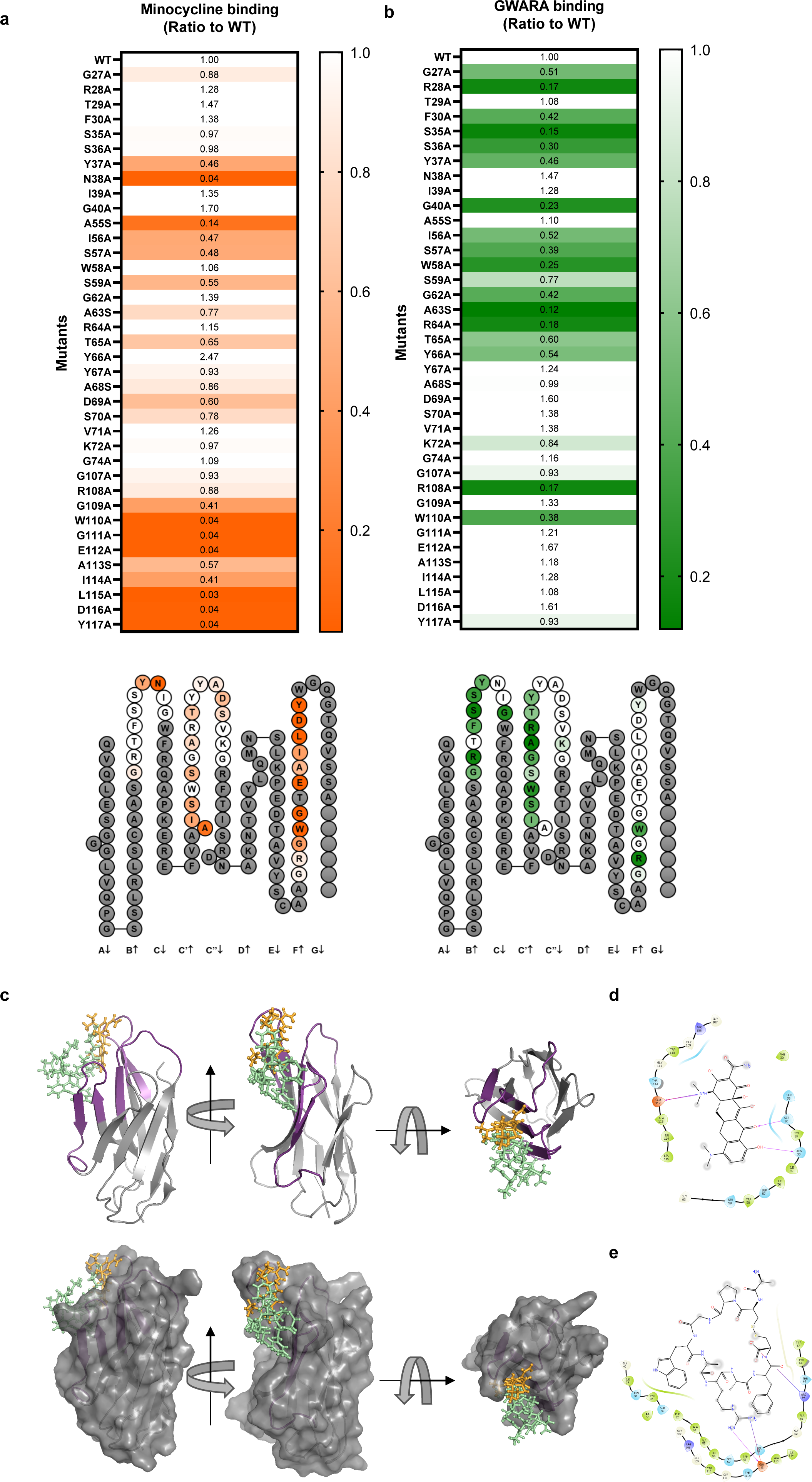
Minocycline and GWARA peptide binding interface for aMinoC1. Alanine scanning was performed on IMGT and Kabat defined CDR regions to maximize the areas of interest: CDR1 positions 27 to 40, CDR2 from 55 to 74 and CDR3 from 107 to 117. Where present, Ala residues were mutated to Serine. **a**, (top) Ratio of minocycline binding kinetics (KD) of wild-type (wt) aMinoC1 and alanine-scan variants (color scale range 0.03-1.0), measured by SPR. Color scale: white for no change (1.0 ratio), dark orange for low binding of variant compared to wt (0.03 ratio). (bottom) Collier de Perles representation of critical amino acid residues (no change:white, lower affinity:orange). Non-mutated residues are shown in grey. **b**, (top) Ratio of binding of wt aMinoC1 and alanine-scan variants to GWARA-Fc peptide by ELISA (color scale range 0.12-1.0). Color scale: white for no change (ratio 1.0), dark green for low binding of variant compared to wt (ratio 0.12). (bottom) Collier de Perles representation of critical amino acid residues (no change:white, lower affinity:green). Non-mutated residues are shown in grey. **c**, Superimposed computational antibody-ligand docking of aMinoC1 with minocycline (orange) and GWARA peptide (green) showing cartoon display (CDR in purple) structure (top) and surface display (bottom) for aMinoC1. **d**, Interaction diagram for aMinoC1 and minocycline. **e**, Interaction diagram for aMinoC1 and GWARA peptide.

Experimental data suggested a disparity of putative contact interactions between antibody/minocycline and antibody/peptide complexes. To determine if a spatial clash was likely to drive the minocycline-peptide competition, we performed computational docking simulations against a 3D model of aMinoC1, using previously defined rotamer conformations for the ligands (Extended Data Fig. 5). Top ranking poses indicated the co-localization of both ligands within the same groove of the antibody (Fig. 3c-e). Despite the limitations of *in silico* modelling, data suggests that steric hindrance is likely to be the main driver of minocycline/peptide competition. Notably, the predicted MHC-I immunogenicity of MinoCAR was low in comparison with widely used clinical components^17,18^ (Extended Data Fig. 6).

### Development and in vitro testing of a novel OFF-switch CAR T cell

We first explored use of this system to generate a controllable CAR. A bipartite CAR architecture (MinoCAR) was constructed consisting of separate antigen recognition and signaling components. The antigen recognition component comprised of a two-arm Fab structure with an anti-EGFR sdAb arm and the aMinoC1 sdAb fused to the transmembrane and endodomain of CD28. The T cell signaling component comprised the GWARA peptide on an extracellular spacer connected to intracellular 41BB and CD3ζ signaling domains. We hypothesized that in the absence of minocycline, these components associate, allowing the CAR to signal in response to antigen, while minocycline would cause dissociation and CAR inhibition (Fig. 4a).

**Figure 4.**
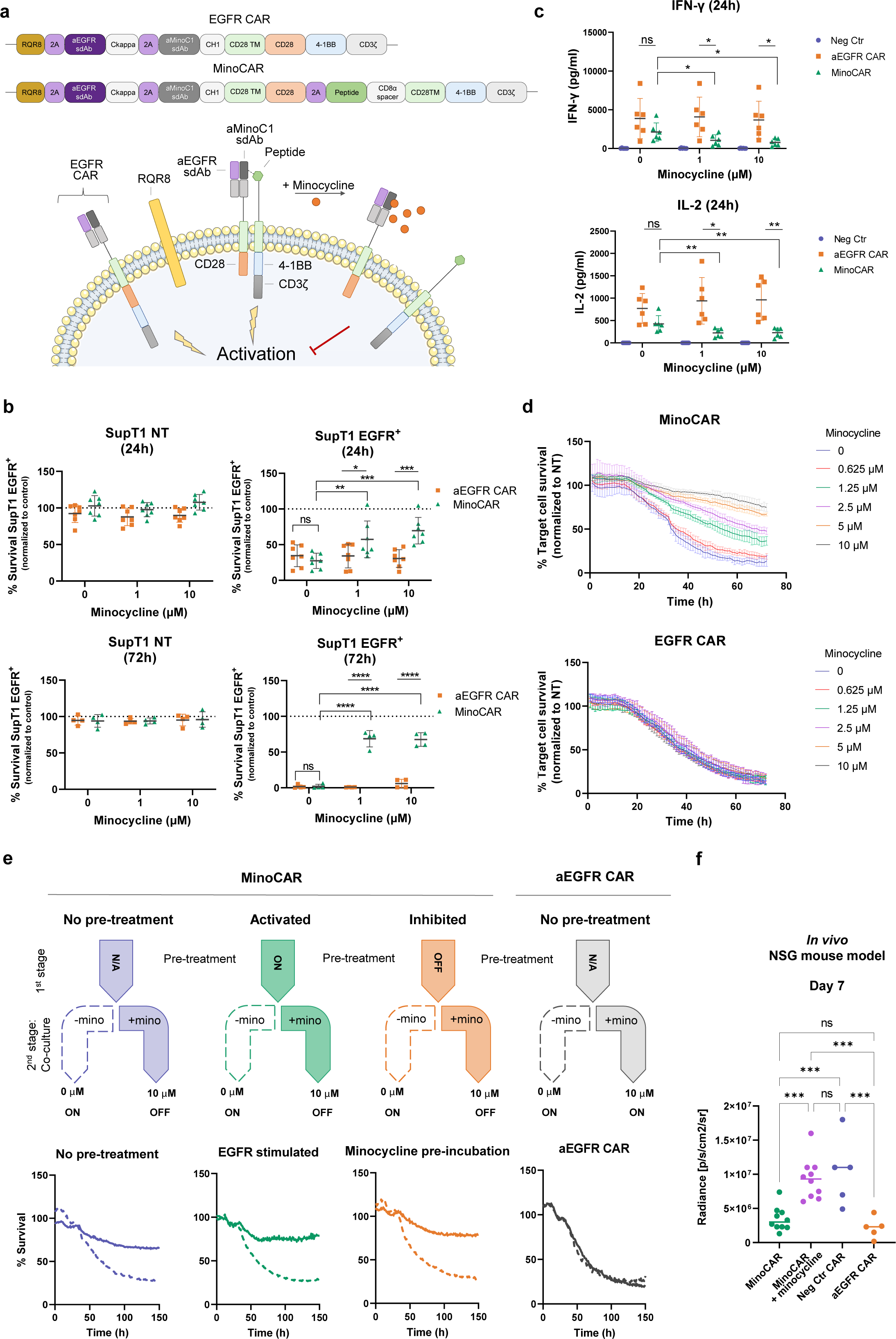
Functional characterization of an OFF-switch CAR T cell (MinoCAR) **a**, Schematic representation of monolithic aEGFR Fab-like CAR (EGFR/aMinoC1 sdAb-TyrpTM-CD28-4-1BBz), and of split MinoCAR (EGFR/aMinoC1 sdAbs-TyrpTM-CD28) with GWARA signaling module (peptide-TyrpTM-4-1BBz). GWARA/aMinoC1 binding inhibited by addition of minocycline. **b**, Cellular cytotoxicity for PBMC transduced with monolithic aEGFR CAR (orange) and MinoCAR (green) against SupT1 NT (left) and SupT1 EGFR^+^ (right) at 24 (top, n=7) and 72h (bottom, n=4), 1:2 E:T ratio. Co-cultures incubated with minocycline at 0, 1 or 10 µM. % survival normalised against NT PBMC. Cells were normalised against negative control MinoCAR carrying an irrelevant peptide (SG_3_S). Mean ± SD, two-way ANOVA with Sidak’s or Tukey’s post-test, * P <0.05, ** P <0.01, *** P <0.001, **** P <0.0001. **c**, INF-γ (top) and IL-2 (bottom) cytokine secretion from PBMC transduced with aEGFR CAR (orange) or aMinoC1 CAR (green) against SupT1 EGFR^+^ target cells (n=6), 1:2 E:T ratio, 24h. Negative Control MinoCAR (blue) carried an irrelevant SG_3_S peptide. Mean ± SD, two-way ANOVA with Dunnett’s post-test, * P <0.05, ** P <0.01. **d**, Kinetics of cytotoxicity of SKOV3 EGFR^+^ target cells co-cultured with PBMC transduced with MinoCAR (green) or aEGFR CAR (orange). Mean ± SD, n=3, 1:2 E:T. Minocycline incubated at a range of concentrations from 0 to 10 µM. **e**, Kinetics of cytotoxicity of SKOV3 EGFR^+^ mKate^+^ co-cultured with PBMC transduced with MinoCAR (left) or aEGFR CAR (right). MinoCAR PBMCs were subjected to no pretreatment (blue), plate-based EGFR stimulation (green) or minocycline incubation (orange). Cells were further incubated with target cells in the presence of 0 µM (dashed arrow) or 10 µM (solid arrow) minocycline. Presence of minocycline in second stage treatment caused inhibition of MinoCAR killing capacity (bottom, solid line) compared to absence of treatment (bottom, dashed line). % survival normalised against NT PBMC. Mean ± SD, n=3, 1:2 E:T ratio. **f**, BLI readout at day 7 post CAR-T injection in a NSG Nalm6 EGFR^+^ tumor mouse model. Significant inhibition of MinoCAR with minocycline injection. One-way ANOVA with Tukey’s post-test, *** P <0.001, ns = not significant.

The cytotoxicity, cytokine release, tunability and reversibility of MinoCAR was tested against both engineered and endogenous EGFR positive cell lines (SupT1 and SKOV-3 respectively) (Extended Data Fig. 7a,b). At 24 and 72 hours, without minocycline, cytotoxicity towards SupT1-EGFR^+^ target cells was observed, equivalent to a control conventional monolithic EGFR-CAR (Fig. 4b). Co-cultures in the presence of minocycline demonstrated a dose-dependent reduction in cytotoxicity and significantly increased target cell survival (average 2.5-fold and 35-fold increase at 24 and 72h, respectively, compared to 0 µM minocycline condition). Similarly, IFN-γ and IL-2 secretion levels in the absence of minocycline were comparable to the conventional EGFR CAR, while minocycline supplementation significantly reduced cytokine secretion (minus 2.7-fold and 1.9-fold for IFN-γ and IL-2, respectively) (Fig. 4c). Notably, MinoCAR was not affected by small molecule analogues doxycycline and tetracycline, or other small molecules such as methotrexate or caffeine (Extended Data Fig. 7c).

### MinoCAR is tunable and can be reversibly controlled

Tunability of MinoCAR was tested by measuring kinetics of cytolysis when exposed to a range of minocycline concentrations. In the absence of minocycline, the rate and extent of cytotoxicity displayed by MinoCAR was comparable to the monolithic EGFR CAR, while increasing concentrations of minocycline induced a dose-dependent decrease in SKOV-3 target cell killing (Fig. 4d). Next, we sought to investigate the ability of MinoCAR T cells to recover activity following inhibition with minocycline and the ability of the drug to inhibit activated cells.

MinoCAR T cells were subjected to three conditions: no pre-treatment, EGFR-induced activation (activated), and minocycline-mediated inactivation (inhibited). Cells were then recovered and cultured with target cells in the presence or absence of minocycline (Fig. 4e). MinoCAR T cells without pre-treatment displayed cytotoxicity in the absence of minocycline and were inhibited by the drug’s presence, as expected. Pre-activated CAR T cells were rapidly inhibited by minocycline and displayed little to no cytotoxicity. Conversely, pre-inhibited CAR T cells quickly recovered activity once the drug was removed, displaying kinetics of target cell killing similar to the “no pre-treatment” condition (Fig. 4e). These results demonstrate that both activation and inhibition of the MinoCAR T cells are rapidly reversible.

In a NOD scid gamma (NSG) Nalm6 EGFR^+^ tumor mouse model, PBMCs transduced with MinoCAR showed significant tumor burden control, comparable to the conventional aEGFR CAR. Mice treated with minocycline showed a significantly reduced MinoCAR efficacy, similar to the effect of a non-functioning CAR (Fig. 4f and Extended Data Fig. 7d).

### sdAb-peptide engager mediated redirecting of cytotoxic T cells

The use of soluble universal receptor engager proteins to direct cytotoxic T cells engineered with a universal CAR has been previously suggested as a therapeutic strategy^19,20^. We sought to explore whether aMinoC1/GWARA could be used to constitute a universal CAR system with additional minocycline control. We hence developed a two-component universal CAR system (MinoUniCAR) consisting of a universal acceptor CAR component comprising the aMinoC1 binding domain, and a soluble functionalizing moiety consisting of a tumor-target binding domain carrying the GWARA peptide tag (Fig. 5a). As a proof of concept, we selected the anti-EGFR sdAb as tumor-targeting adaptor and fused it to the GWARA peptide tag.

**Figure 5.**
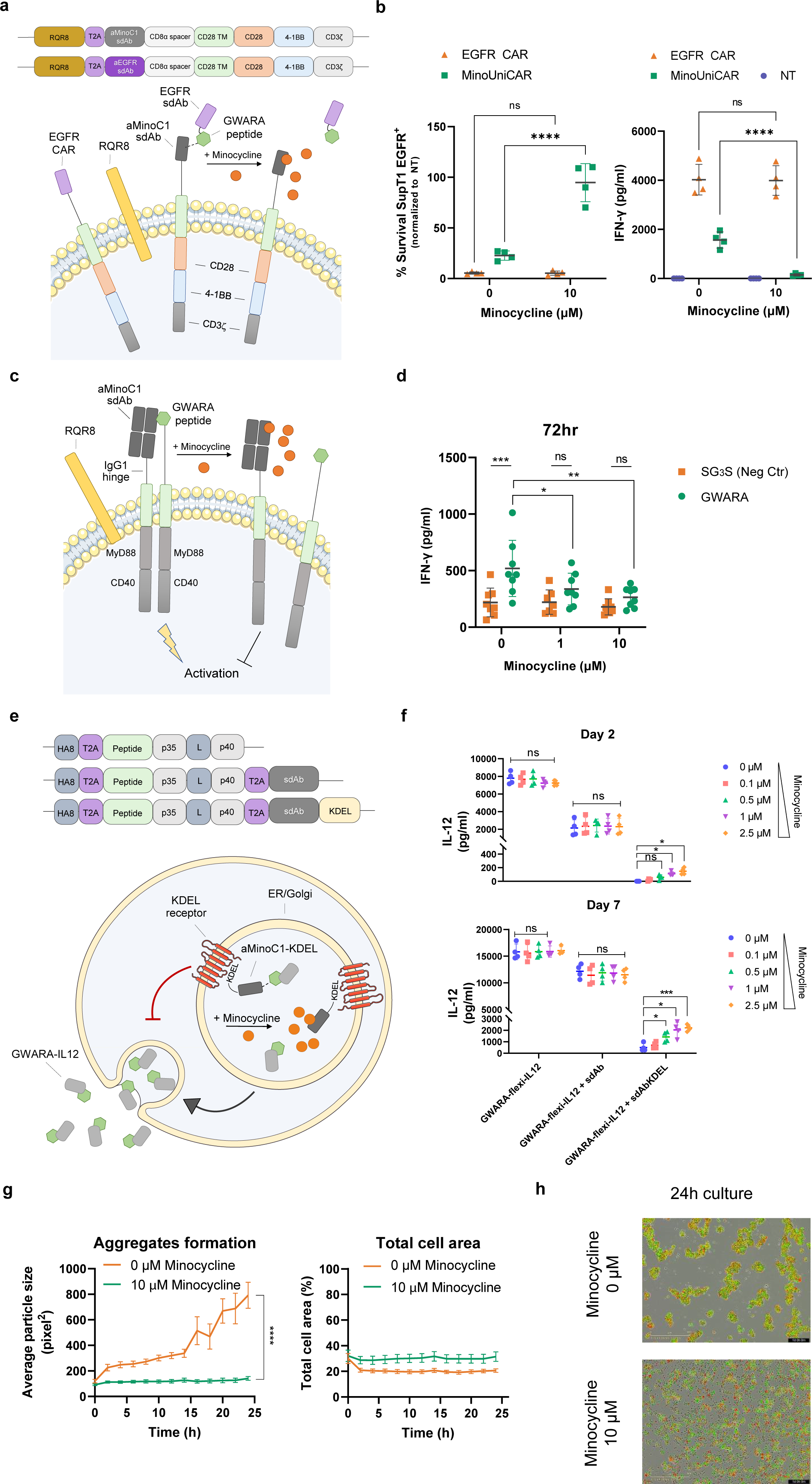
Extended applications for minocycline tuneable control module. **a**, Schematic overview and construct design for aEGFR CAR (aEGFR sdAb-CD8stk-TyrpTM-41BBz), universal adapter CAR MinoUniCAR (aMinoC1-CD8stk-TyrpTM-41BBz) and soluble aEGFR sdAb-GWARA functionalizing molecule. **b**, Cytotoxicity (left) and IFN-γ secretion (right) for transduced PBMC with test CAR constructs (in **a**) co-cultured with SupT1 EGFR^+^ target cells. % target cell survival normalised for non-transduced PBMC. Minocycline incubated at 0 or 10 µM for 24h, 1:2 E:T ratio, n=4. Significant increase in target cell survival and decreased IFN-γ secretion in the presence of minocycline. Two-way ANOVA with Sidak’s post-test, **** P <0.0001. **c**, Schematic representation of controlled signalling module based on MyD88/CD40 endodomains. Construct incudes aMinoC1 Fab-like CAR-IgG1 hinge spacer-TyrpTM-MyD88-CD40. A separate polypeptide carries the GWARA peptide-CD8stk-TyrpTM-MyD88-CD40. Addition of minocycline can dissociate aMinoC1-GWARA binding and prevent signalling. **d**, IFN-γ secretion by PBMC transduced with aMinoC1-Myd88-CD40 construct with GWARA-MyD88/CD40 (green) or control SG_3_S-MyD88/CD40 (orange) constructs. Absence of minocycline showed significant upregulation of IFN-γ by GWARA-MyD88/CD40 compared to SG_3_G-MyD88/CD40. Dose-dependent decrease of IFN-γ secretion visible at increasing concentrations of minocycline. n=8, 72h. Two-way ANOVA with Sikak’s post-test, * P <0.05, ** P <0.01, *** P <0.001. **e**, Schematic of ER/Golgi retention system for minocycline-mediated secretion of GWARA-tagged flexi-IL-12 by aMinoC1-KDEL. Control molecules include untagged aMinoC1 and no aMinoC1. **f**, Minocycline-induced secretion of GWARA-flexi-IL-12 from transduced PBMCs on anti-CD3/anti-CD28 plate-based stimulation. Significant dose-dependent GWARA-flexi-IL-12 secretion at day 2 and 7 for aMinoC1-KDEL in the presence of Minocycline concentrations from 0 to 2.5 µM (n=4). Two-way ANOVA with Dunnett’s post-test, * P <0.05, *** P <0.001. **g**, Time-course cell aggregation formation (left) and total cell area count (right) for SupT1 cells transduced with GWARA-CD8stk-CD28TM and mCherry, co-cultured with SupT1 cells transduced with aMinoC1-CD8stk-CD28TM and eGFP in the presence of 0 (orange) or 10 µM (green) minocycline (n=3). Two-way ANOVA for main effect model, **** P < 0.0001. **h**, Representative snapshot of co-cultures at 24h with 0 µM (top) or 10 µM (bottom) minocycline for SupT1 GWARA-CD8stk-CD28TM and mCherry (red) and aMinoC1-CD8stk-CD28TM and eGFP (green). Co-localisation of signal is visible in yellow.

To test the MinoUniCAR system, the EGFR-peptide adaptor protein was added exogenously to CAR transduced PBMCs at a concentration that was previously determined to provide efficient CAR activation (Extended Data Fig. 8). MinoUniCAR stimulated by EGFR antigen on plate-bound assays showed efficient IFN-y release which was rapidly inhibited in the presence of minocycline with a 10-fold cytokine reduction (148 vs 1569 pg/ml for 10 and 0 µM minocycline, respectively) (Fig. 5b). Similarly, minocycline was able to inhibit the MinoUniCAR cytotoxicity when co-cultured against target SupT1-EGFR^+^ cells, with on average 95% cell survival. In the absence of the drug, the MinoUniCAR recovered cytotoxic capacity. The control EGFR CAR was not affected by minocycline (Fig. 5b).

### Minocycline mediated control of cellular signaling

We next tested whether aMinoC1-GWARA engagement could be harnessed to control cellular signal transmission. As proof of concept, we sought to test a construct which can transmit tunable MyD88/CD40 signals. An inducible MyD88/CD40 inducible system using multimerization induced by FKBP12/AP1097 has been previously described by Foster et al,^21^ and was found to have application in CAR T cell therapy. We generated a dual aMinoC1 Fab architecture fused to an IgG1 hinge, CD28TM and MyD88/CD40 endodomains, paired with a GWARA peptide on a CD8stk with CD28TM and MyD88/CD40 endodomains (Fig. 5c). A version including a SG_3_S peptide instead of GWARA was used as negative control. In the absence of additional stimuli, transduced PBMCs with the aMinoC1/GWARA MyD88/CD40 module showed significantly higher IFN-γ secretion (520 vs 218 pg/ml of negative control), a response that was tuned down by an average of 2-fold, with increasing concentrations of minocycline (Fig.5d).

### Minocycline mediated secretion of an anti-tumor payload

We next investigated whether the minocycline control system could be used to control protein secretion. We hypothesized that fusing the Lys-Asp-Glu-Leu (KDEL) motif to aMinoC1 could promote its retention in the Golgi. Hence, a GWARA peptide-tagged secreted protein would be constitutively directed to the Golgi, while minocycline-mediated dissociation from the aMinoC1/KDEL would allow its secretion.

We used IL12 to test this system. IL12 can potently activity immune responses against cancer but has a narrow therapeutic window with toxicity occurring even when secreted by engineered immune cells. A single chain variant of IL12 (p35 linked to p40, flexi-IL12)^22^ was functionalized with the GWARA peptide (GWARA-flexi-IL12) and expressed alongside aMinoC1 carrying the KDEL retention motif at the C-terminal (Fig. 5e). Control constructs consisted of the GWARA-flexi-IL12 alone or co-expressed with the aMinoC1 sdAb lacking the KDEL motif to prevent retention.

Transduced T cells were monitored at 2- and 7-days post-transduction for cumulative IL12 secretion in the presence of minocycline ranging from 0 to 2.5 µM. ELISA data showed efficient secretion of control GWARA-flexi-IL12 at day 2 and 7 without modulation induced by minocycline (Fig. 5f). Presence of aMinoC1 without KDEL retention caused lower secretory capacity, probably ascribable to a larger transcriptional and translational burden, but similarly unaffected by the presence of minocycline. The GWARA-flexi-IL12 with aMinoC1-KDEL showed no detectable IL12 at day 2 and only 450 pg/ml at day 7. Incremental addition of minocycline triggered a dose-dependent release of IL12 up to 150 pg/ml at day 2 and 2.2 µg/ml at day 7 (Fig. 5f).

### sdAb-peptide engagement as cellular organizers

We hypothesized that the aMinoC1/GWARA interaction could also be used to trigger selective cell-cell interaction and formation of organized cellular aggregates. The suspension cell line SupT1 was engineered to co-express membrane-bound GWARA and mCherry. A second set of SupT1 cells were instead engineered to co-express membrane-bound aMinoC1 and eGFP (Extended Data Fig. 9). Co-culture of the two cell lines with 10 µM minocycline showed a random distribution of cells without clear spatial organization. However, in the absence of minocycline, a time course observation of cells in culture showed the rapid formation of cellular aggregates indicating dynamic cooperation of GWARA and aMinoC1 SupT1 cells (Fig. 5f,g, Extended Data Fig. 9).

## Discussion

Increasingly complex cell-based therapies are being used to treat a range of diseases. Examples include engineered immune cells to treat cancer and autoimmunity,^23,24^ engineered hematopoietic stem cells to treat monogenic disorders^25^ and therapies with iPS-derived cells applied to degenerative diseases^26^. However, unlike small molecule or protein-based therapeutics, cellular therapies may engraft, expand and function in an autonomous fashion. Consequently, therapeutic potency or toxicity cannot easily be controlled by stopping administration or titrating dose. Toxicity can therefore be fulminant and uncontrollable^27^. Control systems which can modulate the activity of engineered cells in response to small molecule pharmaceuticals have been described. These allow “remote control” of cellular therapies and can ensure safety and modulate activity.

Initial control systems exploited the ability of small molecules to induce protein-protein interaction. The earliest designs of such “ON-systems” exploited rapamycin mediated heterodimerization of FKBP12 and the FRB fragment of mTOR^1,28,29^. Wu *et al*. designed a rapamycin controllable CAR by incorporating FKBP12 and FRB components into split antigen recognition and signaling components^4^. Variations of this approach have also been described where FRB and FKBP12 components are extracellular^6^ and where FKBP12-FRB interactions control CAR immune synapse length to tune activity^5^. Additional examples of FKRBP12/FRB use include suicide genes where rapamycin induces Caspase 9 activation^3^. However, rapamycin is immunosuppressive and nephrotoxic^30^ which is a limitation for many applications. To address this, pharmacologically inert “bumped” rapamycin analogs such as AP21967 have been developed which interact with FRB mutated with complementary “holes”, but not with wild-type FRB^31^.

Alternative control systems which also use small molecules to induce protein-protein interaction have been described. These take advantage of pharmacological inhibitors of viral proteases. In one example, an inducible CAR is designed such that a linker recognized by a hepatitis C protease connects antigen recognition and signaling domains of the receptor (SNIP-CAR). The protease is co-expressed resulting in constitutive cleavage with separation of antigen recognition and signaling domains and hence CAR inactivation; small molecule inhibitors of the protease such as grazoprevir and ritonavir prevent this separation, rendering the CAR active^9^.

Alternatively, small molecule control systems can be engineered to disrupt protein-protein interaction (OFF systems). The first example of this was described by Giordano-Attianese et al^11^. Here, the interaction between the mitochondrial protein Bcl-XL and the BH3 domain of BIM was targeted. The authors engineered a human scaffold (LD3) derived from the apolipoprotein 4, with the BH3 motif. The Bcl-XL/LD3 complex could then be displaced by two existing Bcl inhibitors A1331852 and A1155463. Incorporation of the two components into a split CAR resulted in the ability to control CAR activity with the Bcl inhibitors^11^. Similarly, we explored the use of minocycline/tetracycline as a reversible OFF system for managing acute toxicity in the TetCAR system. This consists of a bi-partite split CAR system relying on TetRB and TiP interaction. Displacement by tetracycline/minocycline of the TiP-signaling domain fusion protein from the membrane bound CAR-TetRB portion could inhibit CAR activity in a tunable and reversible manner^12^.

However, several limitations prevent these small molecule control systems from being readily translated into clinical products. Firstly, the use of xenogeneic or unnatural proteins such as the bacterial TetRB^12^ or the viral NS3 protease^9^, or the neoepitopes such as the LD3 and similar engineered scaffolds^11,13^, can trigger immunogenicity^32^. Secondly, designer small molecules such as AP1903 and AP21967 and bcl inhibitors A1331852 and A1155463 have not been granted regulatory approval, greatly hampering clinical use. Finally, in systems where approved small molecules can be used, these small molecules often have significant pharmacologic effects and toxicities (e.g. Rapamycin is a powerful immunosuppressive and is nephrotoxic). The characteristics of an ideal system include minimally immunogenic components which can be controlled by a clinically approved small molecule with little pharmacological effects.

In designing a new system from scratch, we selected minocycline as the control molecule. Minocycline has favorable properties which include excellent biodistribution and a high absorption rate (95-100%). Serum concentrations peak at 2-3 hours and range between 0.7 and 3.9 µg/mL with a 12 to 24 hour half-file in blood^33–35^. Minocycline can cross the blood-brain barrier^36^ and also has excellent tissue penetration, with high tissue/serum concentration ratios in liver and bile (>10) and moderate ratios in several other organs^37^. Notably, minocycline is pharmacologically inert apart from antibiotic properties, allowing for long term use with a good safety profile^38^.

We first sought to isolate a camelid sdAb fragment with specificity towards minocycline. sdAbs are minimally immunogenic due to their high similarity to human VH3-23 family^39^. sdAbs are also attractive as a modular protein control system with their reduced size, enhanced thermal stability and higher hydrophilicity, resulting in lower aggregation propensities^40,41^. The deep paratopes formed by CDR and framework regions of sdAbs, combined with conformational flexibility, enables binding to small molecules (haptens) in a highly specific fashion, with affinity ranges from pM to µM (affinity examples include PP6 dye; 2.5 nM,^42^ picloram; 3-354 µM^43^, auxin; 0.5-20 µM^44^, methotrexate; 29-515 nM^45^, 15-acetyl-deoxynivalenol/15-AcDON; 5-215 µM^46^ and triclocarbon; 0.98-1.37 nM^47^) and with ability to differentiate between similar analogues^48,49^. sdAbs have been employed in the detection of product contaminants^48^, chromatographic extraction^50^, and more recently, for remote controlled biological functions via hapten-induced sdAb dimerization^51^.

We conjugated minocycline to KLH to improve immunogenicity in alpaca. Further, alternating between KLH- and BSA-conjugated minocycline during sequential panning of an immune library ensured specificity towards the hapten rather than linker or carrier protein. The isolated aMinoC1 sdAb displayed strong specificity towards minocycline, in line with previously described antibody-hapten interactions^45–47^, and without cross-reactivity towards the close analogs doxycycline and tetracycline. Additionally, with an affinity of 31 nM it was within the range of affinities previously described for small molecule control systems^11,12^. Binding to minocycline also showed a substantial increase in Tm_50_ (79.5-89.6°C) for the sdAb, suggesting a conformational stabilizing event occurring during binding.

We sought to develop an OFF system. Hence, we required a moiety which would compete with minocycline for sdAb binding. A cyclic 7-mer peptide format was selected as the binding partner, taking advantage of the cysteine-constrained structural integrity to elicit high affinity interactions^52–55^. Phage display panning under stringent competitive elution with minocycline ensured any enriched peptide sequences could be displaceable by the drug. The selected peptide demonstrated an affinity of 111 nM and could be rapidly and reversibly displaced by minocycline.

In the absence of a solved complex structure, due to low resolution crystal formation, we employed an alanine scan approach to investigate how minocycline and the cyclic peptide interacted with sdAb. CDR3 was predominantly involved in minocycline interaction, while CDR1 and CDR2 where required for peptide engagement. This indicates that the dAb interacts with distinct contact points in both cases, however our docking simulations suggest competition for the same groove in the CDR space as the main displacement driver. Interestingly, the aMinoC1 interaction with minocycline and GWARA peptide (KD 31 nM and 111 nM, respectively) closely resembles the TetRB-tetracycline (KD 2.8 nM) and TetRB-TiP peptide (640 nM) interaction^12,56–58^.

We next explored utility of this system to a range of potentially useful applications. We first sought to evaluate the aMinoC1 sdAb system as an OFF-switch split CAR format, showing practical clinical applications for therapeutic modulation. Our data suggested minocycline rapidly displaced the signaling domain, leading to reversible and dose-dependent CAR inhibition by 24 h using a clinically relevant dose of minocycline which is well-tolerated in humans^34^. MinoCAR was comparable to conventional CAR T cells, without significant differences in cytotoxicity or cytokine secretion in the absence of minocycline. A proof-of-concept experiment on NSG mice demonstrated efficient minocycline mediated inhibition of functional MinoCAR T cells *in vivo*. We further developed a universal CAR^59,60^ (MinoUniCAR), demonstrating that the aMinoC1 sdAb/peptide interaction could be harnessed to reversibly functionalize inert CAR T cells and mediate specific anti-tumor activity.

To demonstrate versatility of this system, we also explored other applications. These included an inducible OFF-switch mechanism for constitutively signaling components. AP1097-inducible multimerization MyD88/CD40 systems have been described previously^21,61^. In contrast, our system sought to inhibit constitutive signaling of aMinoC1 and GWARA-peptide fused MyD88/CD40 in the presence of minocycline. Our finding showed comparable levels of IFN-γ secretion to that previously described for two ON-switch MyD88/CD40 constructs^21,61^, with a dose-dependent inhibition mediated by minocycline. Controlled secretion of potent mediators, such as pro-inflammatory cytokines, may be desirable, especially for molecules that are characterized by safety concerns over systemic toxicity^62^. Using IL-12 as an example, we demonstrated controlled secretion by exploiting the ER/Golgi retention signal peptides (KDEL)^63,64^. KDEL-tagged aMinoC1 sdAb could efficiently retain IL-12 (flexi-IL12) when fused to the GWARA peptide. As a final example, we demonstrated that controllable sdAb/peptide interaction could be used to trigger tissue organization by stimulating cell-cell interactions. This system could also be adapted to build customized cell-cell communications for synthetic tissue engineering^65^.

In conclusion, we have developed a novel small molecule control system using minimally immunogenic protein domains and a widely available pharmacologically inert antibiotic as the inducer. We have demonstrated versatility of this system by demonstrating multiple applications. Future improvements may include adapting this system to cytoplasmic applications. In this context, cysteine constraint could be substituted by two anti-parallel coiled coil alpha-helical structures grafted with the GWARA sequence,^66^ using novel linear peptides, or via the generation of a second sdAb component, similar to that described for a caffeine-induced dual sdAb dimerization for transgene expression^67,68^. However, the system in its current form may already have practical utility. Additionally, we hope that this *ab initio* approach to designing control systems around existing pharmaceuticals may be applied to development of multiple novel orthogonal systems.

## METHODS

### Minocycline conjugation

Minocycline was functionalized by introduction of a free sulfhydryl group on a spacer arm to enable maleimide conjugation. Maleimide activated KLH and BSA were used to conjugate the modified minocycline containing the thiol group.

### Immunisation campaign

An Alpaca was immunized using minocycline conjugated to KLH. Following six sub-cutaneous immunizations, sera from the animal was screened to confirm seroconversion against minocycline-BSA via ELISA. Lymphocytes were collected and preserved in RNAlater for the construction of a phage display library.

### Phage Display from immunized Alpacas

Complementary DNA (cDNA) synthesis was carried out using primers designed to amplify the antibody heavy chain-coding region (VHH) from lymphocytes extending from the variable (V) region through to the constant heavy 2 domain (CH2) region. The camelid heavy chain antibody (HCAb) was isolated from classical antibody DNA by agarose gel electrophoresis and further amplified. The double stranded DNA library was ligated into the phage display vector pHEN1 by using unique primers containing SfiI and NotI restriction sites at the 5’ and 3’ends, respectively. The E. coli strain, ER2738, which has a tetracycline resistance gene linked to the F+ gene was transformed via electroporation by incubating ER2738 electrocompetent cells (Lucigen) with ligated DNA in chilled electroporation cuvettes (0.1 cm gap) prior to electroporation using the Bio-rad MicroPulser (EC1 cycle, time constant: 4.5-5.5 ms). An estimated library size of 5×10^8^ unique clones was generated.

The library was panned against biotinylated BSA-conjugated minocycline coupled to streptavidin-coated beads at a concentration of 1 μg/ml. Minocycline-BSA bound phage were eluted using pre-warmed (37°C) Trypsin-EDTA and rotated for 10 minutes at 37°C. 2 selection rounds were carried out, followed by ELISA screening, first analyzing enrichment of the polyclonal library followed by single colony selection, screening and Sanger sequencing.

For ELISA screening, Nunc 96-well plates were coated with minocycline-BSA, minocycline-KLH or BSA only at 1 μg/ml. Plates were blocked with 2% milk in PBS for 1 hour. Whole-Phage, supernatant from soluble dAb and periplasmic extracted dAb was incubated in appropriate wells and incubated for 2 hours at room temperature. Anti-M13-HRP (0.5 μg/ml) was added for the whole phage ELISA and anti-Myc (0.5 μg/ml) was added for soluble protein ELISA and incubated for 1 hour at room temperature.

Analysis of positive ELISA binding data was used to select monoclonal phage for sequencing. PCR amplification of monoclonal phage DNA was performed using a colony PCR reaction. Using the 2x Master Mix OneTaq (M0486L) protocol, PCR reactions containing 1 μl of bacteria from the phage containing supernatant as the template DNA were set up. Amplified DNA was sequenced.

### Phage Display for CX_7_C Peptide Library

Cysteine constrained 7mer (CX_7_C) peptide sequences specific to an aMino sdAb were generated using the Ph.D.™-C7C Phage Display Peptide Library Kit (New England BioLabs, E8120S), a combinatorial library consisting of randomized display peptides with a disulphide constrained loop (AC-XXXXXXX-CGGGS) fused to the pIII coat protein of the M13 phagemid. The library consists of approximately 1 x 10^9^ unique sequences. Phagemid amplification, panning and selection was carried out as previously described with a few methodical exceptions. 3 rounds of panning and enrichment were carried out against biotinylated anti-minocycline single domain antibody clone 1 fused to streptavidin beads. Elution of bound phagemid was carried out using 1 μM minocycline and used directly for subsequent phagemid amplification.

### Differential scanning fluorimetry

The Prometheus NT.48 NanoDSF instrument was used to characterize the thermal and chemical unfolding of aMinoC1 sdAb under native conditions and in the presence of minocycline. A dye-free protocol was used whereby the intrinsic fluorescence of tryptophan and tyrosine was measured by scanning samples at 330 nm and 350 nm to determine protein unfolding. Protein samples were normalized to 0.2-1 mg/ml and supplemented with minocycline at concentrations of 0.0001 μM, 0.001 µM, 0.01 μM, 0.1 μM, 1 μM, 10 μM, 100 μM, 1000 μM and 2500 µM. Melting scan was carried out by setting the run at 1°C/minute temperature increments from 20°C to 95°C. Tm calculated as first derivative of 350/330 nm ratio.

### Surface Plasmon Resonance (SPR)

Surface plasmon resonance (SPR) affinity and kinetic analysis was carried out using the Biacore T200 instrument (GE Healthcare). aMino sdAbs were immobilized to a CM5 sensor chip at a density of 4300-4600 RU. Binding assays were carried out using 1x HBS-EP+ running buffer. Various concentrations (2.5 μM with 2-fold serial dilutions) of the analyte were injected for 150 second at 30 μl/min with 150 second dissociation time. Glycine-HCl (pH 2.0) was used as the regeneration buffer for the sensor chips. For the Alanine scanning experiments, the anti-minocycline antibody (sdAb-muIgG2aFc fusion) was captured on a Protein A series S sensor chip, to a density of 3000 RU. Minocycline was injected at a concentration of 5 μM with 2-fold serial dilutions, with 150 second contact time and 300 second dissociation at 30 μl/minute. Glycine-HCl pH 1.5 was used as regeneration buffer. In each case, flow cell 1 used for reference subtraction and a ‘0 concentration’ sensogram of buffer alone was used as a double reference subtraction to factor for drift. Data analysis was carried out using the Biacore T200 Evaluation Software, version 3.0 and Biacore Insight evaluation software. The 1:1 Langmuir binding model was used to calculate the association (ka), dissociation (kd) rate constants and the equilibrium dissociation constant (KD).

### Isothermal Titration Calorimetry (ITC)

ITC measurements were performed using the PEAQ-ITC non-automated (MicroCal) at 25°C. Antibody sample was dialyzed overnight using 5 L of PBS (20 mM Na₃PO₄, 150 mM NaCl, pH 7.4). Minocycline (Sigma-Aldrich) was dissolved in DMSO, followed by dilution in PBS such that the final DMSO concentration was 1%. The concentration of the anti-minocycline sdAb was determined using extinction coefficients ɛ280nm= 31,065 M-1cm-1 The single domain antibody (2 μM) in the cell was titrated with minocycline (50 μM) using 22 injections of 10 μl made at 120 second intervals with a stirring speed of 300 rpm. The binding isotherm plot was fitted by non-linear regression using the Origin software to a one set of sites 1:1 binding model to generate the thermodynamic parameters of the antibody-minocycline interaction.

### Expression and purification of proteins

Antibodies were expressed by transient transfection in Expi-CHO cells as murine IgG2a Fc domain conjugates and purified using the HiTrap MabSelect SuRe (GE Healthcare) affinity chromatography. Briefly, MabSelect SuRe 1 ml column (GE Healthcare) was equilibrated with 5 column volumes of PBS pH 7.4 at a flow rate of 1 ml/min. Supernatant was applied to the column using Akta™ Pure system at a flow rate of 1 ml/min. The column was then washed with 10 column volumes of PBS pH 7.4 at 1 ml/min. Samples were eluted from the column using 3 ml of IgG elution buffer (Pierce, 21004) at 1 ml/min and directly loaded through a double stacked HiTrap 5 ml desalting column. Samples were collected on a 96-well plate using a fraction collector unit at a fraction volume of 250 μl. Fractions were analyzed using SDS-PAGE to confirm presence of appropriate size protein band and purity of protein sample. LCAR-B38M antibody sequences were obtained from patent literature^69,70^. aEGFR VHH antibody sequence was obtained from the literature^71^.

### Cell lines

HEK-293T cells (ATCC; ATCCCRL-11268) and SKOV-3 cells (ATCC; ATCC HTB-77) were cultured in Iscove’s modified Dulbecco’s medium (IMDM) supplemented with 10% FBS (Labtech) and 2 mM GlutaMAX (Invitrogen). SupT1 cells (ECACC; 95013123) were cultured in complete RPMI (RPMI-1640, Lonza) supplemented with 10% FBS and 2 mM GlutaMAX. SupT1 cells were genetically modified by transduction with an SFG vector to express human EGFR. ExpiCHO cells were cultured in ExpiCHO medium (Gibco) using Erlenmeyer shake flasks (Corning) and maintained in a Kuhner shaker at 37°C and 8% CO_2_, at 225 rpm.

### Transduction

γ-retroviral supernatants were produced by transiently transfecting HEK-293T cells (3 × 10^6^) with an RD114 envelope expression plasmid (a gift from M. Collins, UCL), and a Gag-pol expression plasmid (a gift from E. Vanin, Baylor College of Medicine), and SFG transgene plasmid. The transfection was carried out using GeneJuice (Millipore) in accordance with manufacturer’s guidelines.

Blood was obtained from buffy coats purchased from NHSBT (NC07). PBMCs were isolated from buffy coats via density gradient sedimentation using Ficoll. PBMCs were activated using anti-CD3 antibody and anti-CD28 antibody (Miltenyi Biotec). 24 hr post activation, culturing media was supplemented with 100 IU IL-2 (2BScientific Limited). At 72 hr, 1 × 10^6^ PBMCs were plated on retronectin coated 6-well plates (Takara Clonetech) with retroviral vectors and centrifuged at 1,000 × *g* for 40 min. 72 hr post transduction, transduction efficiency was assessed and PBMCs were maintained in complete RPMI medium supplemented with 100 IU IL-2.

### Flow cytometry

Flow cytometry was performed using the MACSQuant® Analyzer 10 or MACSQuant® X (Miltenyi Biotec). All flow cytometry data was analyzed using FlowJo v.7.6.2 software (Tree Star Inc, Ashland, OR). Cell staining was carried out by incubation with PBS containing the recommended concentration of antibodies at room temperature for 30 minutes. PBS washes were carried out between antibodies staining. Cells viability was determined using viability dye SYTOX™ Blue Dead Cell Stain (ThermoFisher) prior to Flow cytometric analysis. Cells were first gated for singlet population identified by FSC-H and FSC-A. Next, live cells were identified using a viability dye. Followed by gating for target cell populations. The antibodies used in the study are as follows: CD3 PE-Cy7 (Biolegend, 317334), Human CD34 APC-conjugated antibody (R&D system, FAB7227A), Human CD34 Alexa Fluor® 488-conjugated antibody, R&D system, FAB7227G), Streptavidin PE, Biolegend, 405204), APC anti-human EGFR antibody (Biolegend, 352905), Alexa Fluor® 488 anti-human EGFR antibody (Biolegend, 352907), PE anti-HA.11 Epitope Tag antibody (Biolegend, 901517), Anti-M13-HRP (Sino Biologics, 1197-mm05T-H), Anti-Myc-HRP (Genscript, A00863) and SYTOX™ Blue Dead Cell Stain (ThermoFisher, S10274).

### FACS-based killing assays

Transduced CAR T cells were determined by staining for the transduction marker RQR8 and normalized by addition of non-transduced T cells. Effector cells were co-cultured with 5.0 × 10^4^ number of target cells (SupT1-NT, SupT1-EGFR+, SKOV3-mKate) cells to achieve desired effector cell to target ratio. Where appropriate, minocycline was supplemented as a part of the culture conditions. Co-cultures were incubated for 24-72 hours at 37°C and 5% CO_2_. After incubation, plates were centrifuged at 400 *g* for 5 minutes, and the supernatant containing secreted cytokines was collected for cytokine analysis. Cells were stained with anti-hCD34-APC and anti-CD3-PeCy7 to differentiate effector T cells and target cells. Cells were washed with 300 μl PBS and stained with SYTOX™ Blue Dead Cell Stain dye. The percentage of target cell survival was measured relative to the number of live target cells co-cultured with non-specific CAR T cells or non-transduced PBMCs.

### Reversibility and tuneability assays

EGFR+ SKOV3-mKate were seeded at 1.0 × 10^4^ cells per well in a TC-treated flat bottom 96-well plate. For reversibility experiments, MinoCAR ON-OFF kinetics were tested by pre-activating CAR T cells by incubating cells for 2 hours on plates pre-coated with recombinant EGFR (10 μg/ml). CAR T cells were washed using 15 ml of PBS prior to co-culture. MinoCAR OFF-ON kinetics were tested by first inhibiting CAR T cells by incubating cells in RPMI supplemented with 10% FBS, 1% GlutaMAX and 10mM minocycline. Cells were then washed using 15 ml of PBS prior to co-culture. Untreated CAR T cells were used as a control and were directly preceded to co-culturing set. In each respective well, 0.5 × 10^4^ transduced CAR T cells were co-cultured with EGFR+SKOV3-mKate target cells with and without minocycline (10 μM). The Incucyte® ZOOM Live-Cell Analysis System was used to carry out image capture at a rate of one image every 1hr for 150 hours. Tuneablity experiments were carried out co-culturing transduced PBMCs expressing the MinoCAR and EGFR-CAR (positive control) with target cells at an effector to target ratio of 1:2 and in the absence and presence of minocycline (0.625 μM, 1.25 μM, 2.5 μM, 5 μM, 10 μM). Image capture rate of each condition was set at 1 image per hour for 72 hours.

### Cytokine-release assays

IFN-γ, IL-2 and IL-12 secretion was measured by collecting supernatant from the respective cell-based assay and frozen at −20°C prior to analysis by ELISA. IFN-γ, IL-2 and IL-12 ELISAs were carried out using the Human IFN-γ ELISA MAX Deluxe kit, Human IL-2 ELISA MAX Deluxe kit and Human IL-12 (p70) ELISA MAX Deluxe kit in accordance with the manufacturer’s instructions (BioLegend).

### sdAb-peptide cytotoxic T cell engager assays

Transduced PBMCs were stained using for RQR8 expression and V5 markers. Cells were then normalized to 50% efficiency by diluting with non-transduced T cells. The tumor targeting adaptor protein (EGFR sdAb) fused to the GWARA peptide was separately expressed by CHO cell transient transfection and purified using His-tag chromatography. For the assay set up, 0.5 x 10^4^ effector cells were co-cultured with SupT1-EGFR+ target cells at a ratio of 1:2 in the presence of 15 μg/ml adaptor protein, with and without minocycline supplementation (10 μM) for 24 hours. IFN-γ secretion was measured as described above.

### IL-12 secretion assays

Transduced PBMCs were stained for the independent HA expression marker using an anti-HA-PE antibody to determine transduction efficiency. Cells were normalized to 60% using non-transduced T cells prior to assay set up. For the assay set up, 0.5 x 10^4^ transduced cells were cultured in RPMI (RPMI-1640) supplemented with 10% Fetal Bovine Serum (FBS), 2 mM GlutaMAX and IL-2 at a concentration of 100 UI/ml. Culture media was also supplemented with and without minocycline supplementation (0.1 μM, 0.5 μM, 1 μM and 2.5 μM). The supernatant was collected at day 1 and day 7. IL-12 ELISA was carried out following manufacturer’s instructions (BioLegend, 431704).

### Cell-cell binding assays

SupT1 cells were transduced to express GWARA-CD8a spacer-CD28TM-2A-mCherry and aMinoC1 sdAb-CD8 spacer-CD28TM-2A-eGFP. 0.5 x 10^4^ cells were co-cultured in RPMI (RPMI-1640) supplemented with 10% Fetal Bovine Serum (FBS) and 2 mM GlutaMAX. Where appropriate, culture media was supplemented with minocycline (10 μM). Images were captured using the IncuCyte real-time imager. Time course images were analyzed using ImageJ (NIH v1.53t) using image analyzer (20 px^2^-infinite size, 0.0-1.0 circularity).

### Homology modelling

Homology model generation and antibody-ligand docking was carried out using Schrodinger BioLuminate software. Camelid single domain antibody templates were selected from the Protein Data Bank (PDB). The optimal template was selected as an appropriate template framework based on the composite scores, structural identities and PDB resolutions. In total, 10 predicted CDR3 loop were generated. The quality of the homology models and amino acid backbone conformations were assessed using Ramachandran plots. The Schrodinger Protein Preparation Wizard PrimeMini Package was utilized to reduce rigidity and optimize the homology model via energy minimization.

Alanine scan mutagenesis of the CDR regions, as defined by the IMGT and Kabat numbering system, was carried out to generate 38 mutant anti-minocycline sdAbs independent alanine mutations or serine mutations where alanine was substituted. Critical hotspot residues were determined using Biacore SPR binding data to determine KD values for the interaction of minocycline or ELISA binding data to determine binding to GWARA peptide. Computational antibody-ligand docking was carried out by preparation of ligands using the Schrodinger LigPrep suite to convert 2D ligand (minocycline or GWARA) structures to produce corresponding low energy 3D structures. A receptor grid was placed on the anti-minocycline sdAb model for ligand docking, centered on the critical residues defined by SPR and ELISA. The GlideScore function of the Schrodinger BioLuminate Glide suite was used to rank the docking models. In combination with experimental binding data from the alanine mutants, top poses were selected for analysis.

### Xenograft model with NALM6

All animal studies were carried out in accordance with a UK Home Office–approved project license and were approved by the UCL Biological Services Ethical Review Committee. NSG mice (female, aged 6–10 weeks) were acquired from Charles River Laboratories and raised under pathogen-free conditions. On day −4, 0.5 × 10^6^ Nalm6 cells engineered to express EGFR and HA-luciferase were injected intravenously into NSG mice. Tumor engraftment was assessed through bioluminescent imaging, employing the IVIS Spectrum system (PerkinElmer) following intra-peritoneal injection of 2 mg D-luciferin. Photon emission from EGFR^+^ NALM6-FLuc cells, expressed in photon per second per cm2 per steradian, was quantified using Living Image software (PerkinElmer). On day 0, mice were randomly assigned to different cohorts prior to intravenous injection of 1 × 10^6^ CAR T cells. The experiment was conducted without blinding; however, the use of bioluminescent imaging provides an objective assessment of tumor growth in this model. For mice treated with minocycline, a stock solution of 4 mg/ml minocycline hydrochloride was prepared by reconstituting minocycline hydrochloride (Sigma) in sterile PBS. Intra-peritoneal injection of 100 μl (0.4 mg) were administered every 2-3 days. Mice were regularly weighed every 2 days, mice exhibiting any of weight loss exceeding 10%, signs of graft-versus-host disease, or disease progression were humanely euthanized.

### Statistical analyses

GraphPad Prism 9.0 (Graphpad Software Inc., La Jolla, CA, RRID:SCR_000306) was used to carry out statistical analyses and calculation. Specific statistical tool utilized is describe in figure legend where appropriate. Statistically significant differences were determined when the p values were < 0.05. All data is presented as mean ± SD unless otherwise stated.

## Acknowledgement

Cell graphics were drawn by using pictures from Servier Medical Art. Servier Medical Art by Servier is licensed under a Creative Commons Attribution 3.0 Unported License.

## Author contributions

M.P. conceived the project and helped design experiments. R.J., A.K. designed and carried out the experiments, analyzed the data and wrote the manuscript. R.B., A.B., T.I., S.O., M.F. generated and characterized the binders and the peptides. R.B., I.G. performed the *in vitro* assays. C.A., K.L., A.D., F.P., J.S. carried out the molecular cloning. A.H. carried out the *in vivo* work. S.T., M.F., M.P. supervised the project, reviewed the data, and wrote the manuscript.

## Ethics declaration

R. Jha, A. Kinna, A. Bulek, I. Gannon, C. Allen, K. Lamb, A. Dolor, F. Parekh, J. Sillibourne, S. Thomas, M. Ferrari and M. Pule are employees and hold equity in Autolus Limited. R. Bughda, T. Ilca, S. Cordoba and S. Onuoha are former employees and may hold equities in Autolus Limited.

Patent applications on the work described in this paper have or may be filed by Autolus Limited.

## Data and materials availability

All data is available in the manuscript or the supplementary information.

