## Supplementary material for "Designer small molecule control system based on Minocycline induced disruption of protein-protein interaction"

**Extended Data Fig. 1. Chemical conjugation of minocycline**

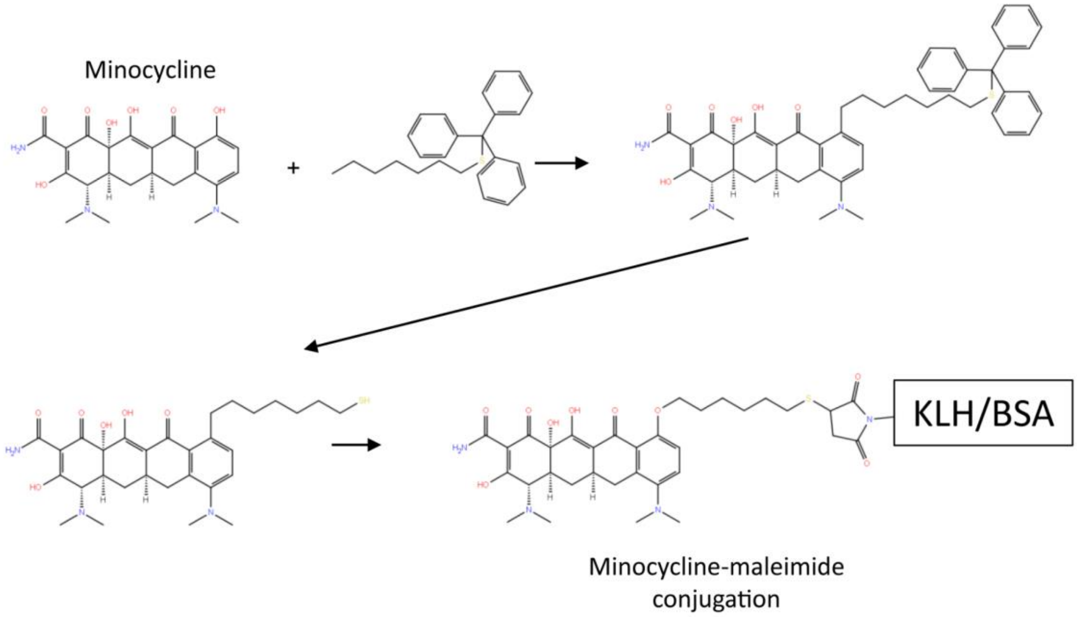


Functionalization of minocycline by addition of a free sulfhydryl group on a spacer arm to enable maleimide conjugation to either (KLH) or (BSA) for immunization and subsequent panning strategies, respectively.

**Extended Data Fig. 2. Alpaca immunization and phage display output**

**a**

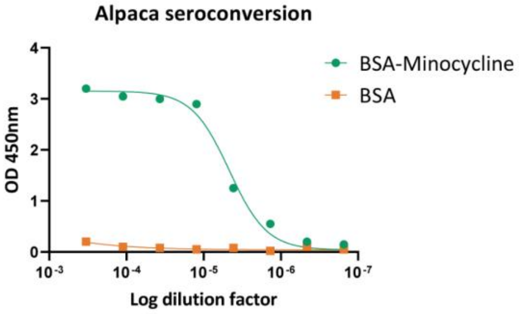


**b**

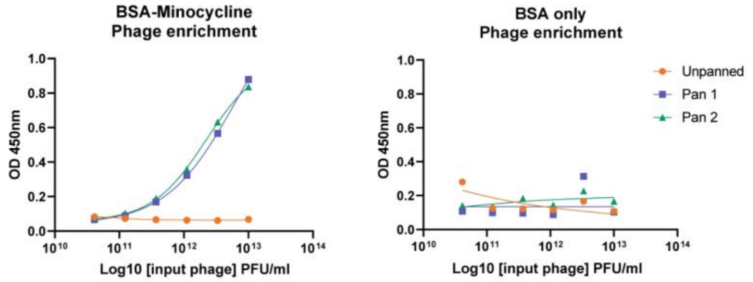


**c**

| Clone | CDR 1 | CDR 2 | CDR 3 |
|---|---|---|---|
| **aMinoC1** | GRTFSSYN | ISWSGART | AAGRGWGTEAILDY |
| **aMinoC4** | GRSLSSYV | ISWSGART | AAGRGWGTEAILDY |
| **aMinoC9** | GRTFSSYN | ISWSGART | VAGRGWGTEAILDY |
| **aMinoC14** | GRTFSNYN | INWSGGRT | AAGRGWGTEAILDY |
| **aMinoC21** | GRTFSSYN | ISWSGART | AAGRGWGTEAILDY |
| **aMinoC22** | GRTFSRYN | ISWSGART | AAGRGWGTEAILDY |
| **aMinoC25** | GRTFSSYN | ISWSGART | AAGRGWGTEAILDY |
| **aMinoC26** | GRTFSSYN | ISRSGGIT | AAGRGWGVEAILDY |
| **aMinoC31** | GNIGLVSV | ITGGGST | RLVNNGRPF |
| **aMinoC32** | GRLSLSSYV | ISWSGART | AAGRGWGTEAILDY |
| **aMinoC36** | GRTFSNYN | INWSGGRT | AAGRGWGTEAILDY |
| **aMinoC37** | GRTFSSYN | ISWSGART | AAGRGWGTEAILDY |
| **aMinoC38** | GRTFSSYN | ISWSGART | AAGRGWGTEAILDY |
| **aMinoC40** | GRTFSSYN | ISWSGART | AAGRGWGTEAILDY |

**d**

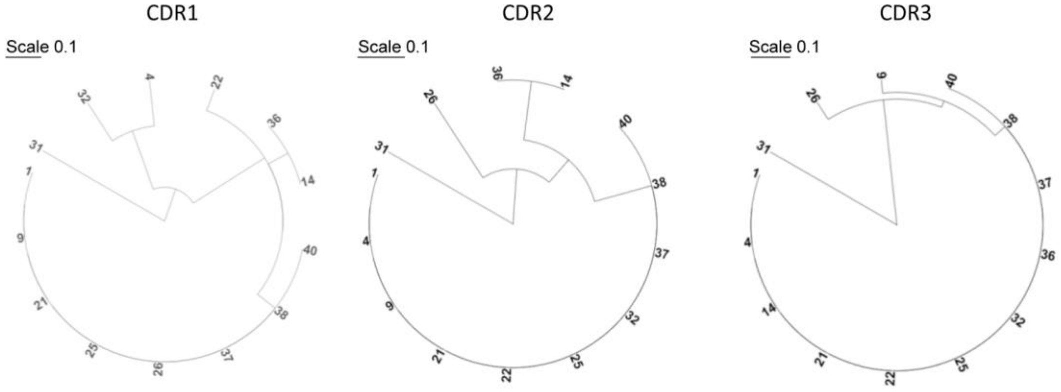

**a**, Immunised Alpaca serum showing specific antibody response to minocycline in ELISA against immobilised BSA-minocycline (green) or BSA only (orange), detected with anti-alpaca IgG-HRP conjugated secondary antibody. **b**, Minocycline specific phage enrichment ELISA for Pan 0 (unpanned, orange), Pan 1 (blue) and Pan 2 (green) against BSA-minocycline (left) and BSA only (right). Bound phage clones detected with anti-M13-HRP conjugated secondary antibody. **c**, Complementarity-determining region (CDR) amino acid sequence analysis of phage derived anti-minocycline sdAb antibodies. **d**, Phylogenetic divergence of CDR1, CDR2 and CDR3 in selected anti-minocycline clones.

**Extended Data Fig. 3. SPR kinetic affinity of anti-minocycline sdAb**

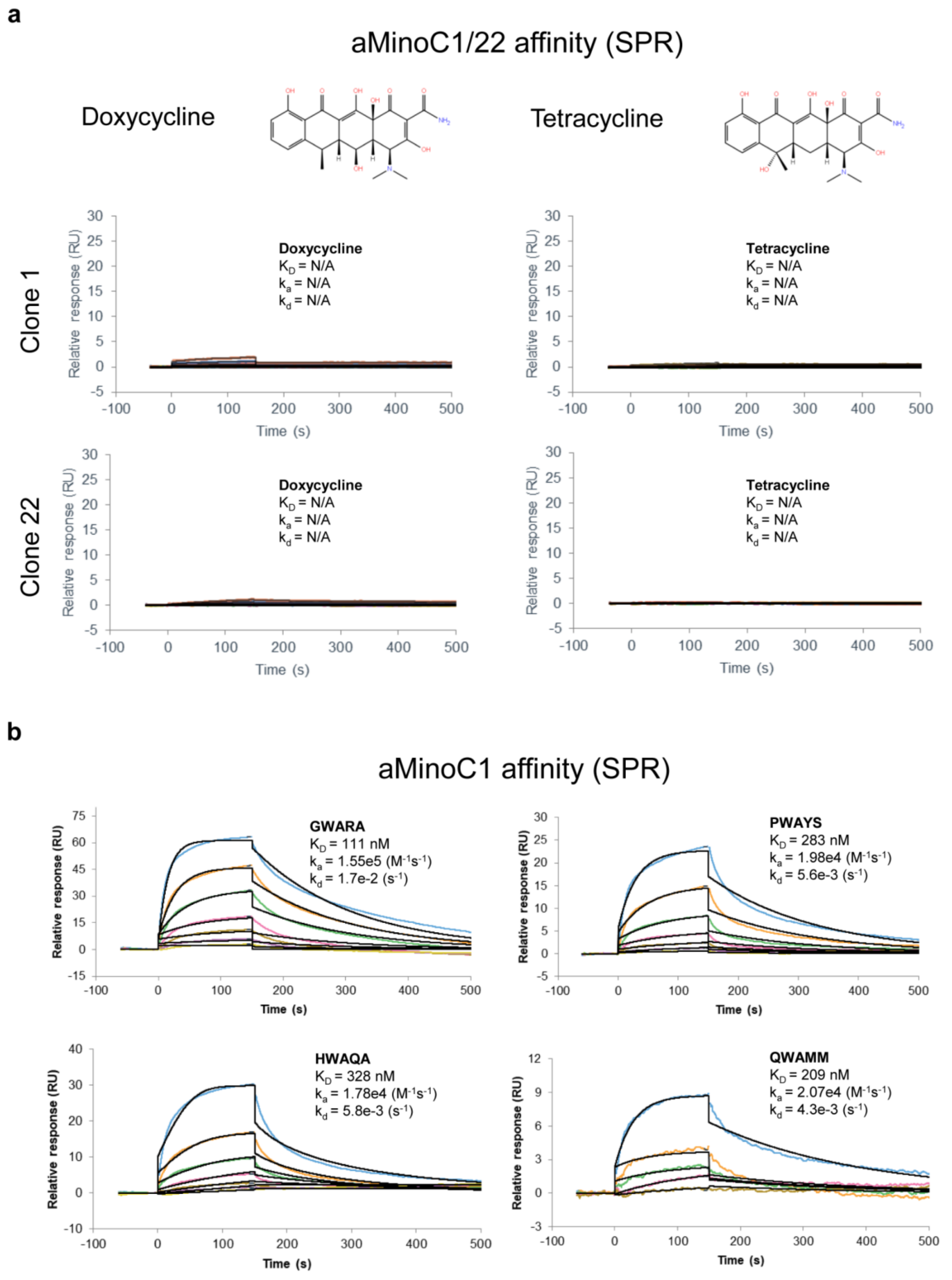


**a**, Surface plasmon resonance (SPR) of aMinoC1 and aMinoC22 sdAb-Fc clones against doxycycline (left) and tetracycline (right) showing lack of binding to structurally related molecules to minocycline. **b**, SPR kinetic affinity of GWARA, PWAYS, HWAQA and QWAMM peptide-Fc against aMinoC1. Kinetics fit with a 1:1 Langmuir binding model. Affinities were measured at 111 nM (GWARA), 283 nM (PWAYS), 328 nM (HWAQA) and 209 nM (QWAMM).

**Extended Data Fig. 4. Dynamic binding of aMinoC1-CX$_7$C peptide-minocycline**

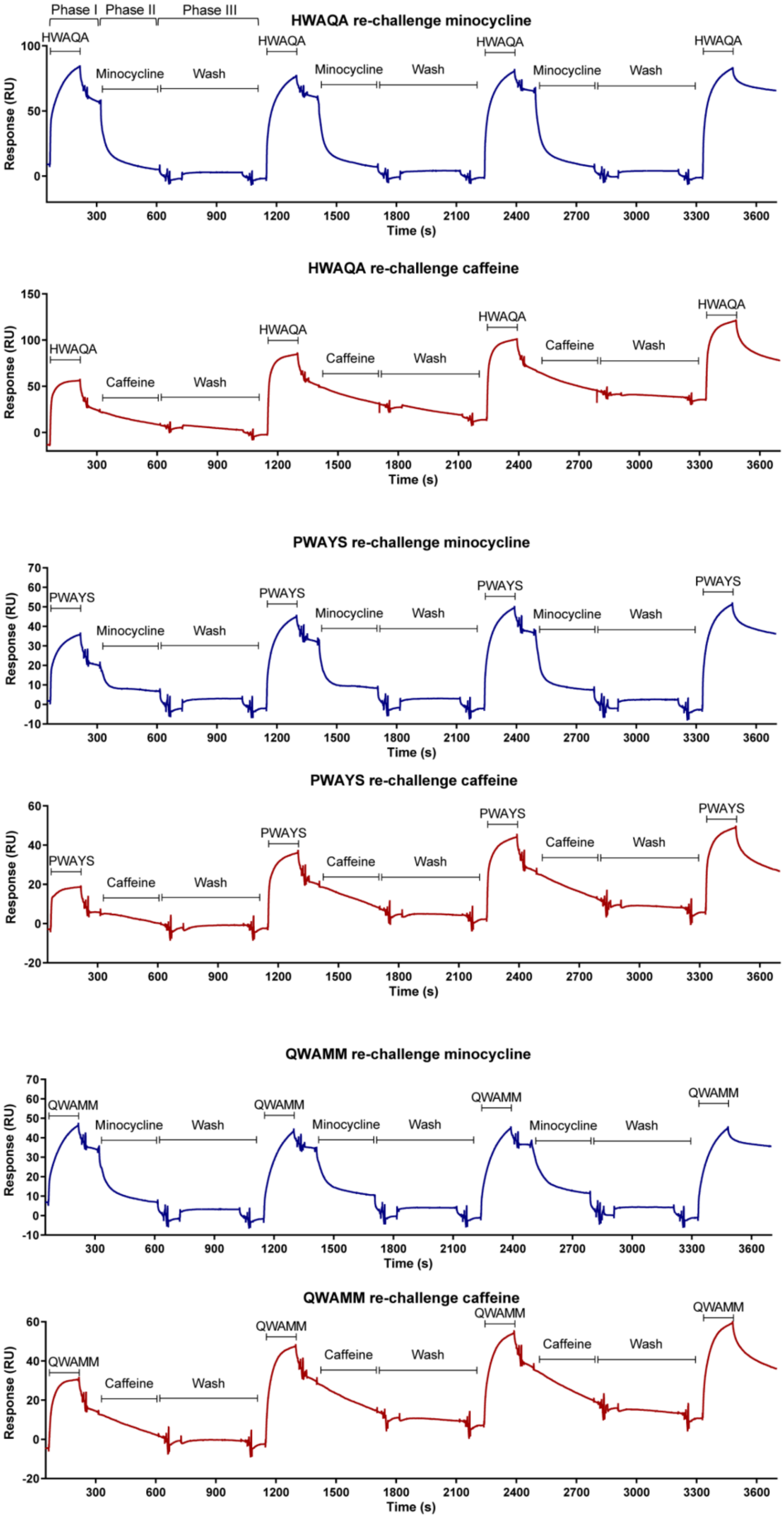

Dynamic minocycline (blue) or caffeine (red) small molecules and peptide-Fc (HWAQA, PWAYS and QWAMM) binding to immobilised aMinoC1 sdAb on Biacore T200. Sequential injections of GWARA-Fc (Phase I), small molecule (Phase II) and dissociation (buffer) step (Phase III) showing minocycline-driven acceleration of peptide-Fc dissociation. Serial challenges with peptide and small molecule show reversibility of the system. No enhanced dissociation visible with caffeine injections.

**Extended Data Fig. 5. Minocycline ligand preparation**

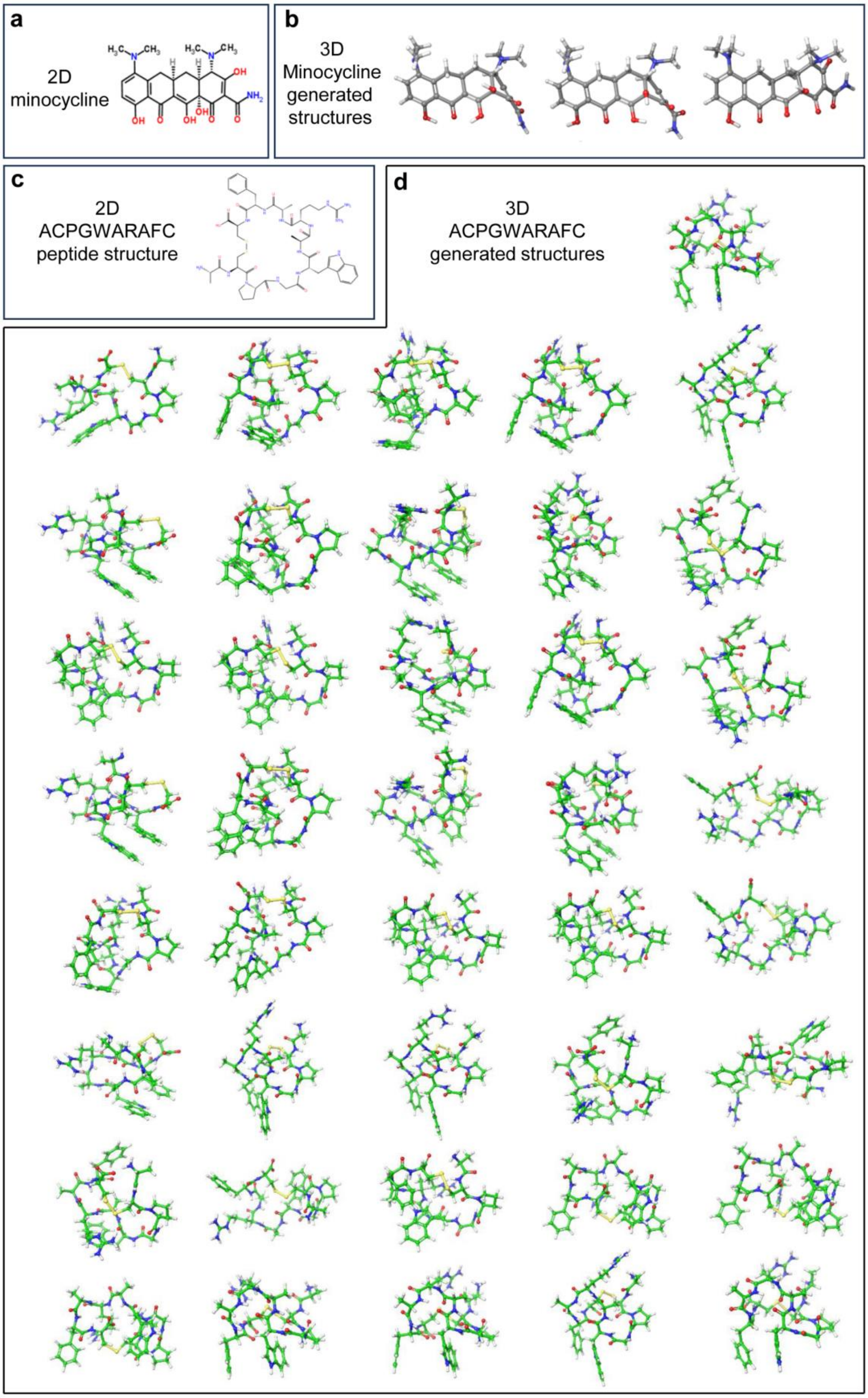

**a**, 2D minocycline structure. **b**, Energy minimised 3D structures for minocycline generated by Schrodinger LigPrep suite. **c**, 2D cyclic GWARA (ACPGWARAFC) peptide structure. **d**, Energy minimised 3D structures for GWARA peptide generated by Schrodinger LigPrep suite.

**Extended Data Fig. 6. *In silico* immunogenicity analysis**

a

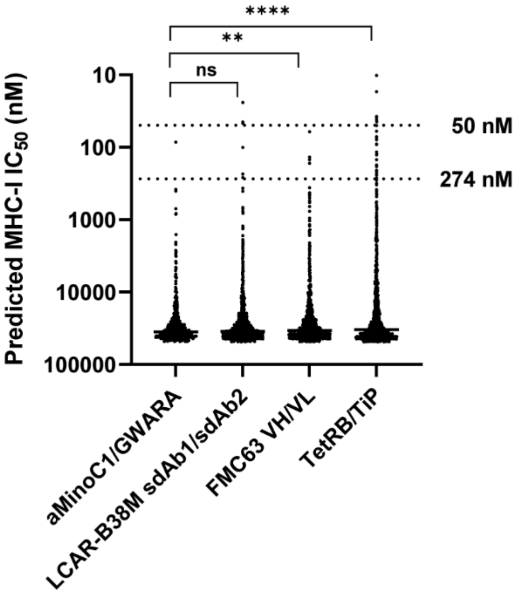


b

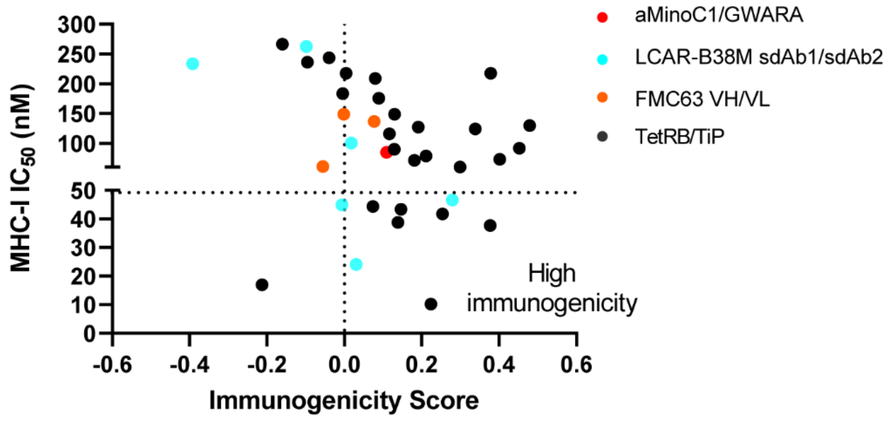


**a**, Predicted MHC binding affinity for 9-14mer peptides derived from aMinoC1 and GWARA peptide, TetRB/TiP, FMC63 VH/VL and LCAR-B38M sdAbs. Affinity calculated using IEDB prediction tool. Dotted lines indicate cut-off for IC50 affinities of 274 nM (threshold for MHC-I engagement) and 50 nM (threshold for immune activation). Significantly lower IC50 affinities for aMinoC1/GWARA compared to bacterial TetRB/TiP and murine FMC53 VH/VL. One-way ANOVA with Dunnett's post-test, ** P <0.01, **** P <0.0001. **b**, MHC-I $IC_{50}$ and immunogenicity score for 9-14mer peptides from aMinoC1/GWARA (red), LCAR-B38M (teal), FMC63 VH/VL (orange) and TetRB/TiP (black) with < 274 nM MHC-I $IC_{50}$ (from **a**). Positive score denotes potential immunogenicity. Peptides with < 50 nM MHC-I $IC_{50}$ and positive immunogenicity score are considered highly immunogenic. No highly immunogenic peptides predicted for aMinoC1/GWARA.

**Extended Data Fig. 7. Characterization of EGFR+ cell lines and MinoCAR cytotoxicity**

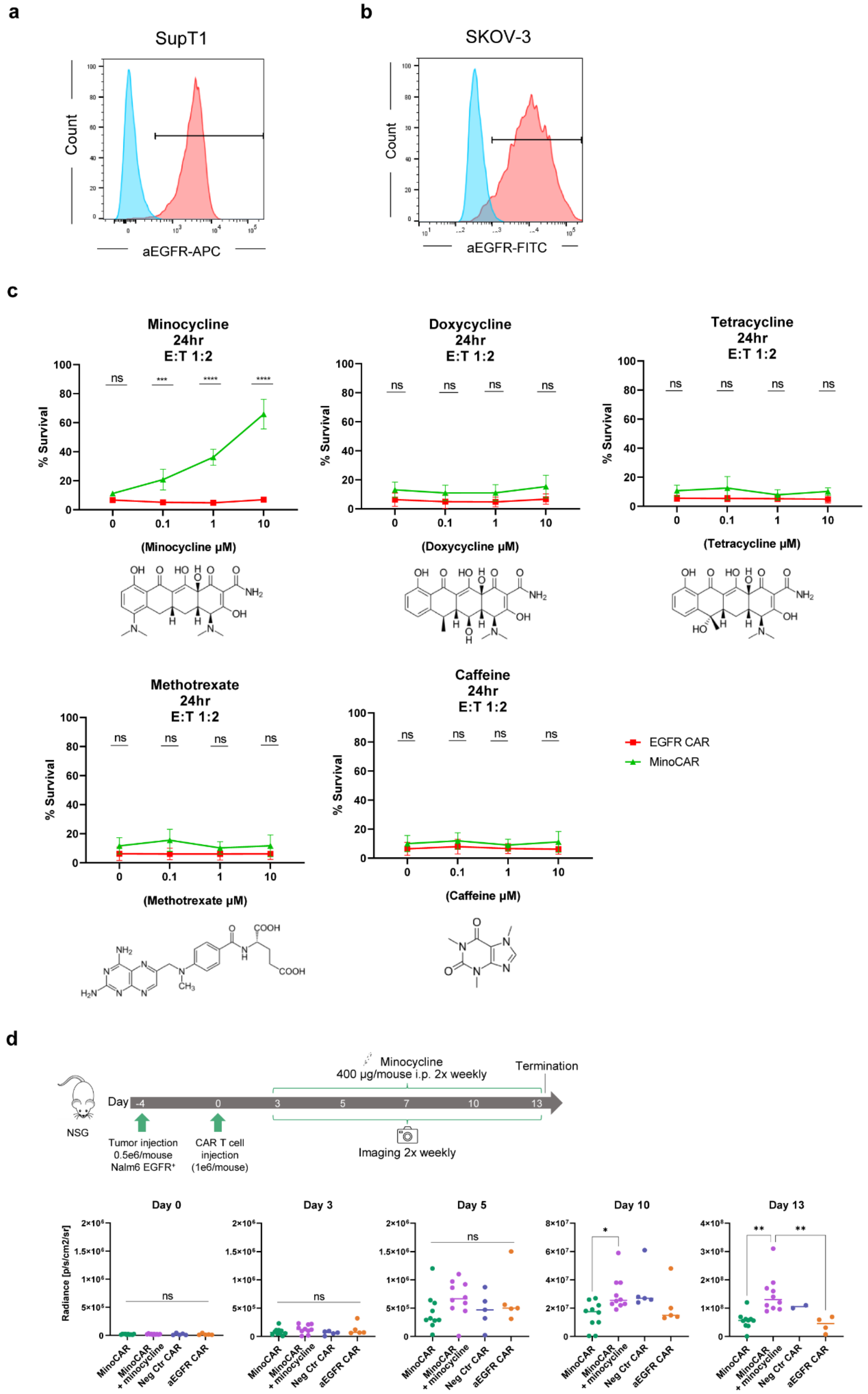

**a**, SupT1 (blue) and SupT1 EGFR$^+$ cells (red) cells stained with anti-EGFR-APC antibody to confirm expression of EGFR marker. Histogram plot of cell count vs EGFR expression (MFI). **b**, SKOV-3 mKate+ cells stained with anti-EGFR-APC to detect natively expressed EGFR (red). Unstained cells presented in blue. Histogram plot of cell count vs EGFR expression (MFI). **c**, Flow cytometry-based cytotoxicity assay of PBMCs transduced with MinoCAR (green) and a conventional anti-EGFR CAR positive control (EGFR-CAR, red). Sup-T1 cells expressing EGFR (SupT1-EGFR+) were used as target cells. Effector cells were co-cultured with target cells at a ratio of 1:2 with varying concentrations of the indicated drug (0.1 µM, 1 µM and 10 µM) for 24 hours. The percent of live target cells was normalized to negative control. Gating strategy was as follows: Singlets (FSH/FSA)>Live cells (SYTOX blue)>Anti-CD3-PeCy7. Data represented as mean ± SD, n=4, two-way ANOVA with Sidak's post-test, *** P <0.001, **** P <0.0001. **d**, (Top) Schematic of NSG Nalm6 EGFR+ tumor mouse model treated with MinoCAR (n=10), aEGFR CAR (n=5) or a negative control CAR carrying the $SG_3S$ peptide (n=5). An additional cohort of MinoCAR included 400 µg/mouse twice weekly i.p. injections of minocycline (n=10). (Bottom) BLI readouts for CAR treated cohorts at days post car injection (DPI) 0, 3, 5, 10 and 13. DPI 7 is reported in Fig. 4f. A significant difference in tumor burden control was detected for MinoCAR and MinoCAR + minocycline cohorts at DPI 7-13. One-way ANOVA with Tukey's post-test, * P < 0.05, ** P <0.01, ns = not significant.

**Extended Data Fig. 8. Dose response of GWARA-EGFR sdAb adaptor to UniMinoCAR**

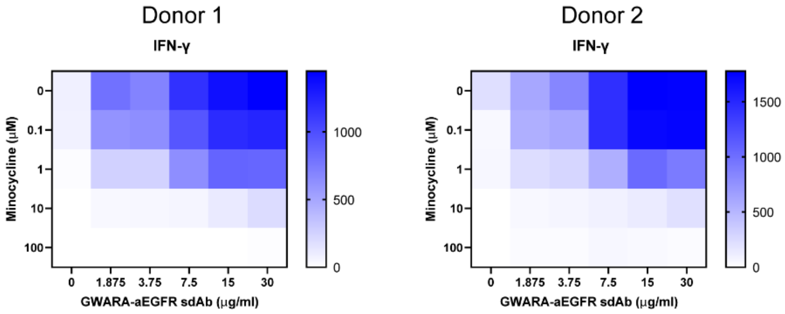


Dose response titration for GWARA-EGFR sdAb adaptor (from 0 to 30 μg/ml) in the presence of 0 to 100 μM minocycline. IFN-γ secretion measured from EGFR plate-based stimulation of transduced PBMCs with MinoUniCAR construct in each matrix condition. Two independent donors analysed.

**Extended Data Fig. 9. Induced cell-cell interaction with aMinoC1/GWARA module**

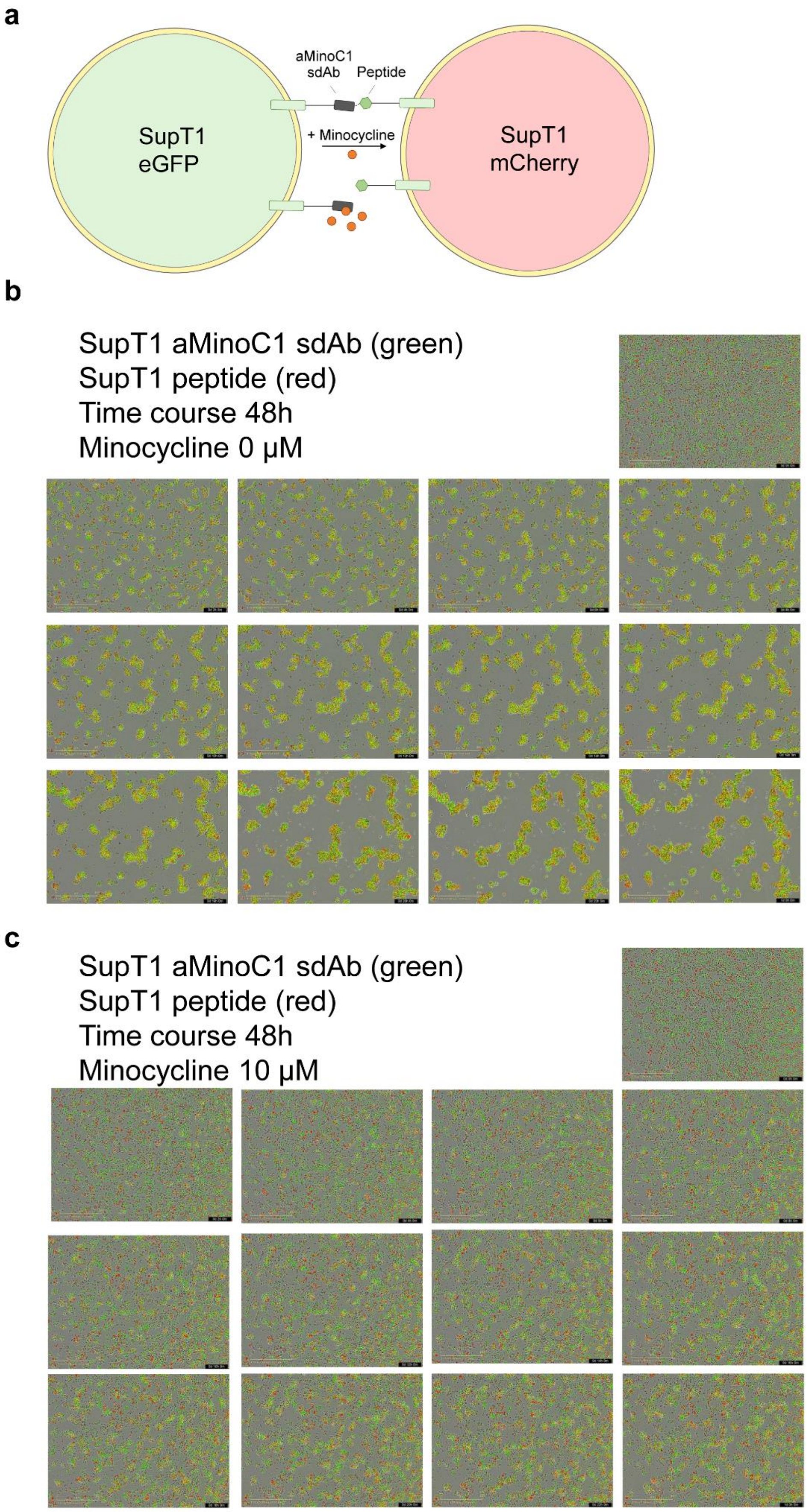

**a**, schematic representation of SupT1 cells transduced with GWARA-CD8stk-CD28TM and mCherry (red) and SupT1 cells transduced with aMinoC1-CD8stk-CD28TM and eGFP (green). **b**, Time course co-cultures of SupT1 GWARA-CD8stk-CD28TM mCherry (red) and SupT1 with aMinoC1-CD8stk-CD28TM eGFP (green) for 24h in the absence of minocycline. Formation of aggregates is visible during co-culture. **c**, Time course co-cultures of SupT1 GWARA-CD8stk-CD28TM mCherry (red) and SupT1 with aMinoC1-CD8stk-CD28TM eGFP (green) for 24h in the presence of 10 μM minocycline. Images acquired with IncuCyte at 2h intervals, representative of 3 independent experiments.
